# Melastatin subfamily Transient Receptor Potential channels support spermatogenesis in planarian flatworms

**DOI:** 10.1101/2024.09.01.610670

**Authors:** Haley Nicole Curry, Roger Huynh, Labib Rouhana

**Affiliations:** Department of Biological Sciences, Wright State University, 3640 Colonel Glenn Hwy., Dayton, OH 45435, USA; Department of Biology, University of Massachusetts Boston, 100 William T. Morrissey Blvd., Boston, MA 02125-3393, USA

**Keywords:** Spermatogenesis, planarian, Transient Receptor Potential Channel Melastatin, TRPM

## Abstract

The Transient Receptor Potential superfamily of proteins (TRPs) form cation channels that are abundant in animal sensory systems. Amongst TRPs, the Melastatin-related subfamily (TRPMs) is composed of members that respond to temperature, pH, sex hormones, and various other stimuli. Some TRPMs exhibit enriched expression in gonads of vertebrate and invertebrate species, but their contributions to germline development remain to be determined. We identified twenty-one potential TRPMs in the planarian flatworm *Schmidtea mediterranea* and analyzed their anatomical distribution of expression by whole-mount *in situ* hybridization. Enriched expression of two TRPMs (*Smed-TRPM-c* and *Smed-TRPM-l*) was detected in testis, whereas eight TRPM genes had detectable expression in patterns representative of neuronal and/or sensory cell types. Functional analysis of TRPM homologs by RNA-interference (RNAi) revealed that disruption of *Smed-TRPM-c* expression results in reduced sperm development, indicating a role for this receptor in supporting spermatogenesis. *Smed-TRPM-l* RNAi did not result in a detectable phenotype, but it increased sperm development deficiencies when combined with *Smed-TRPM-c* RNAi. Fluorescence *in situ* hybridization revealed expression of *Smed-TRPM-c* in early spermatogenic cells within testes, suggesting cell-autonomous regulatory functions in germ cells for this gene. In addition, *Smed-TRPM-c* RNAi resulted in reduced numbers of presumptive germline stem cell clusters in asexual planarians, suggesting that *Smed-TRPM-c* supports establishment, maintenance, and/or expansion of spermatogonial germline stem cells. While further research is needed to identify the factors that trigger Smed-TRPM-c activity, these findings reveal one of few known examples for TRPM function in direct regulation of sperm development.

## INTRODUCTION

Studies of animal reproduction are relevant for preservation of natural resources, raising livestock, forestry, fishery, and protection of crops from pests. Environmental influences, such as temperature, are known to alter endogenous endocrine pathways that regulate seasonal reproduction either directly by serving as a cue for activation developmental processes, or indirectly by altering metabolic pathways that influence fecundity (Chmura and Williams, 2022). As climate change continues to threaten the survival of hundreds of species, our need to understand how organisms sense changes in environmental conditions, as well as the molecular mechanisms behind their influence on mating, fertility, and fecundity, is of outmost urgency.

Planarian flatworms are free-living members of the superphylum Lophotrochozoa that are best known for their regenerative abilities. Although fission and regeneration serve as means for asexual reproduction of some planarians, most species reproduce sexually as cross-fertilizing hermaphrodites (Hyman, 1951, Vila-Farré ant Rink, 2018; Issigonis and Newmark, 2019). Planarians that live in habitats with fluctuating temperatures have been reported to switch reproductive modes seasonally (Kenk, 1935). Sexualization can also be induced experimentally in the laboratory (Kobayashi et al., 2004), and the signaling molecules that ignite or modulate sexual maturation are starting to be identified (Collins et al., 2010; Kobayashi et al., 2017; Rouhana et al., 2017; Nakagawa et al., 2018; Issigonis et al., 2023). Furthermore, reproductive structures regress in response to injury or starvation, and redevelop upon healing and growth, showing extreme developmental plasticity (Newmark et al., 2008).

Given their dynamic reproductive biology, planarians provide a great opportunity to learn about the molecular mechanisms behind external influence on sexual reproduction. The sexual organs and entire reproductive system of planarians, including the germline, develop post-embryonically from somatic pluripotent stem cells called neoblasts (Kobayashi et al., 1999; Wang et al., 2007). Planarians also continuously replenish, grow, and regenerate their entire anatomy through differentiation of neoblasts (Newmark and Sanchez Alvarado, 2002). Given that expression of genes can be disrupted systemically by supplementing food with double-stranded RNA (dsRNA) of target-specific sequence (Rouhana et al., 2013), interruption of continuous cellular turnover allows the discovery of genes required for development of tissues (including the germline). Systemic RNAi in planarians also allows for identification of genes required during sexual maturation and development of reproductive structures without interfering with embryonic development. This approach has uncovered novel genes required for germline development, as well as previously unknown roles in gametogenesis for genes characterized in other systems.

This study includes characterization of genes predicted to encode Transient Receptor Potential (TRP) channels of the Melastatin subfamily (TRPM) in the planarian *Schmidtea mediterranea*. TRPMs mediate cellular responses to environmental and physiological cues that include hormones, changes in pH, and temperature (Diver et al., 2022; Zheng, 2013). Mammalian TRPMs that respond to changes in temperature are expressed in somatosensory neurons, where they play roles in noxious and innocuous temperature response (Kashio and Tominaga, 2022). However, some members of the TRPM subfamily are also expressed during mammalian spermatogenesis, where they are thought to influence sperm development and function (Martínez-Lopez et al., 2011; De Blas et al., 2009; Lee et al., 2003; Borowiec et al., 2016). One of the TRPM homologs in *S. mediterranea*, *Smed-TRPM-c*, displays testis-specific expression and is required for spermatogenesis, providing an avenue to study the contributions of TRPMs to regulation of sperm development in planarians. In asexual planarians, *Smed-TRPM-c* is required for normal maintenance of presumptive germline stem cells, suggesting a function in the earlier stages of spermatogenesis. Altogether, these results complement findings in mice to indicate that TRPMs may have ancestral role during spermatogenesis and suggest that TRPMs may regulate spermatogenesis in response to changes in temperature.

## RESULTS

### Identification of Transient Receptor Potential Melastatin subfamily channels in *Schmidtea mediterranea*

To identify putative TRPMs in the planarian flatworm *S. mediterranea*, TBLASTN searches for homologs of human TRPM3 (NCBI GenBank ID NP_066003.3) were performed against transcriptomes of sexual *S. mediterranea* deposited in PlanMine (the planarian flatworm sequence repository; Brandl et al., 2016; Rozanski et al., 2013). Hits were further analyzed by reciprocal BLAST searches against annotated human proteins. Twenty-one non-redundant putative TRPM genes were identified (Supplementary Table S1). Analysis of identical sequences by BLASTN in reference transcriptomes of the asexual biotype of *S. mediterranea* revealed matches for all 21 putative TRPM genes (Supplementary Table S1), indicating that these are expressed at levels detectable by RNAseq in asexual planarians. Three of these genes are orthologs of TRPMs previously characterized in the planarian *Dugesia japonica* (Inoue et al., 2014). The genes represented by contigs dd_Smed_v6_17857_0_1 and dd_Smed_v6_26481_0_1 on PlanMine are orthologs of DjTRPMa and were therefore named *Smed-TRPM-a1* and *Smed-TRPM-a2*, respectively (BLASTP E-values = 0.0 and 2.04 × 10^−135^). The gene represented by contig dd_Smed_v6_9288_0_1 was named *Smed-TRPM-b*, as it is orthologous to DjTRPMb (E-value = 0.0). The remaining 18 TRPM homologs in *S. mediterranea* were named *Smed-TRPM-c* to *-t*. The sequence of *Smed-TRPM-a* to *-e*, and four additional genes, more closely matched TRPM3 than any other human TRPM in reciprocal BLAST results, while others more closely matched human TRPM1, 2, 4, 5, 6, and 8 (Supplementary Table S1). These results indicate an expansion of TRPM genes to at least 21 homologs in *S. mediterranea*.

### Smed-TRPM-c and Smed-TRPM-l are expressed in testis lobes of S. mediterranea

To determine the anatomical distribution of expression of planarian TRPM genes in *S. mediterranea*, whole-mount *in situ* hybridization (WMISH) analyses were performed using sexually mature planarian hermaphrodites. Using riboprobes that corresponded to ∼500 bps of reference cDNA contig sequence, *Smed-TRPM-a1* expression was detected in cells concentrated at the head tip and further distributed throughout the planarian body (Figure 1A), matching the expression of its ortholog *DjTRPMa* (Inoue et al., 2014) in thermosensory neurons. *Smed-TRPM-a2* expression was only detected in the gut (Figure 1B). Expression of *Smed-TRPM-b*, -*f*, *-h*, and *-m* was detected in the brain region (Figure 1C, 1G, 1I, 1K, and 1N), with WMISH signal for some of these genes being limited to subsets of putative neurons (Figure 1G’ and 1N’) or very faint (Figure 1C’ and 1I’). *Smed-TRPM-c* expression was only detected in testis lobes, which are abundantly located from the tail to the region posterior to the head along the dorsolateral anatomy of *S. mediterranea* (Figure 1D and D’). Expression of *Smed-TRPM-j* and *-n* were observed along the periphery of the head tip (Figure 1K and 1O), whereas *Smed-TRPM-r* expression was enriched in subsets of cells that reside along the ventral nerve cords and brain (Figure 1S to S’’). *Smed-TRPM-e*, *-f*, *-o*, and -*s* expression was detected through much of the gut (Figure 1F, 1G, 1P, and 1S), while expression of *Smed-TRPM-l* was detected in the gut (Figure 1M) and testis (Figure 1M’). *Smed-TRPM-o* was detected in the gut and pharynx (Figure 1P). It is important noting that the planarian gut and copulatory apparatus tend to generate background signal in WMISH analyses using colorimetric development, therefore some of the signals from these structures were deemed inconclusive (Supplementary Table S1). The TRPM homologs with no detected or inconclusive expression include *Smed-TRPM-d*, *-g, -i*, *-k*, *-p*, *-q*, and *-t* (Figure 1E, 1H, 1J, 1L, 1Q, 1R, and 1U).

**Figure 1.**
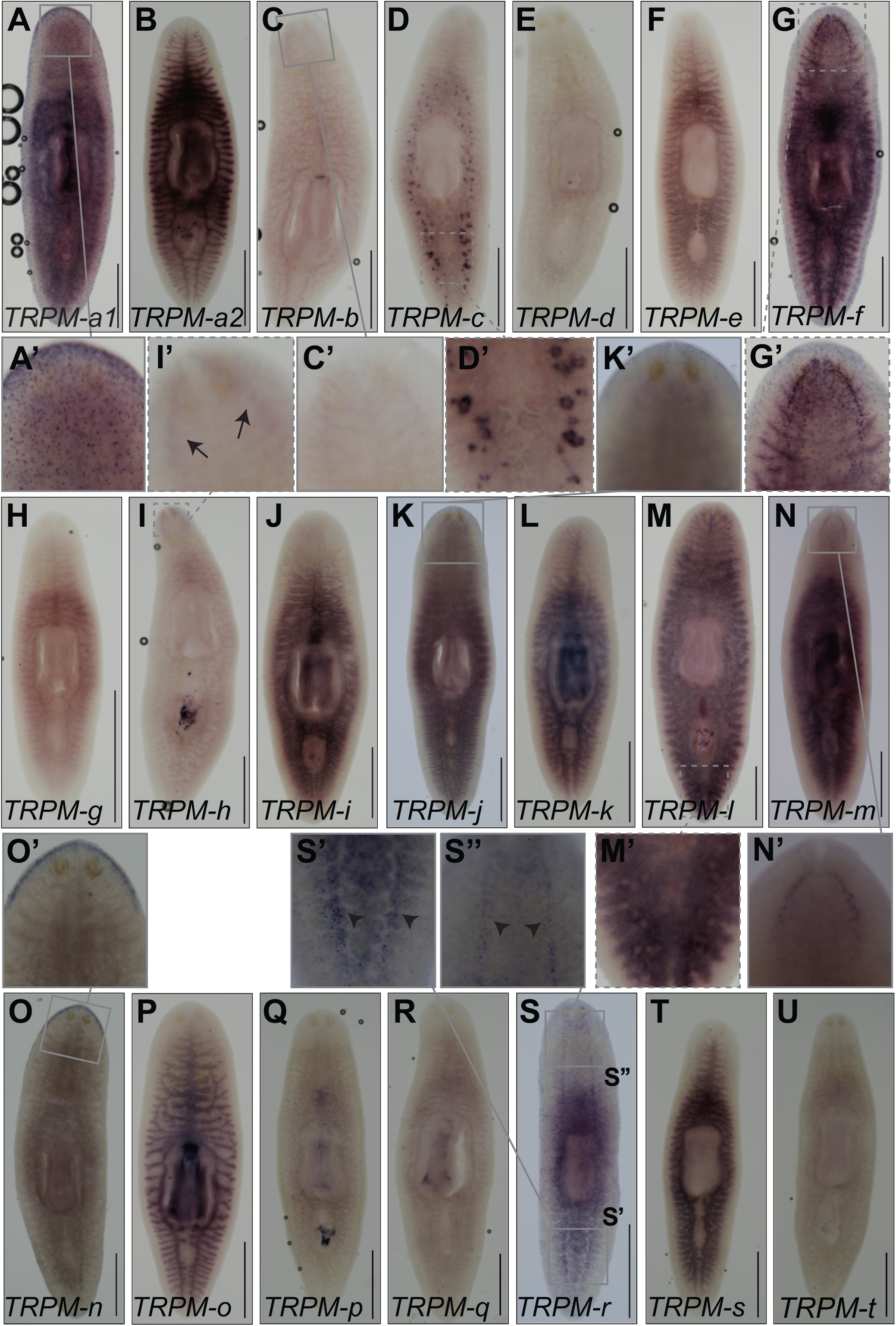
Analysis of expression of TRPMs in *S. mediterranea* reveals two homologs expressed in testes. **(A-U)** Whole-mount colorimetric *in situ* hybridization analyses using partial sequence riboprobes (∼500 nts) display distribution of expression of *Smed-TRPM-a1* (*TRPM-a1*; A), *Smed-TRPM-a2* (*TRPM-a2*; B), and *Smed-TRPM-b* to *-t* (*TRPM-b* to *-t*; C-U). Insets show magnified view of planarian testis lobes in the posterior dorsal region of planarians (D’ and M’), as well as sensory cells (A’ and G’), brain region (C’, G’, I’, N’), head periphery (K’ and O’), and cells along the posterior (S’) and anterior (S”) ventral nerve cords. Arrows point at the location of brain lobes in (I’), and arrow heads point at posterior ventral nerve cords in (S’ and S”). Scale bars = 1 mm.

### *Smed-TRPM-c* is required to maintain normal levels of sperm development

To test whether TRPMs expressed in planarian testes are required for sperm development, juvenile (sexually immature) planarian hermaphrodites were subjected to continuous RNAi treatments in groups targeting either *Smed-TRPM-c* (*Smed-TRPM-c(RNAi)*), *Smed-TRPM-l* (*Smed-TRPM-l(RNAi)*), or both *Smed-TRPM-c* and *Smed-TRPM-l* (*Smed-TRPM-c;TRPM-l(RNAi)*), by mixing dsRNA corresponding to ∼500 bps of gene-specific sequence in their food. A control group was fed dsRNA with sequence that corresponds to firefly *Luciferase*, which does not interfere with planarian homeostasis or germline development (Magley and Rouhana, 2019; Lesko and Rouhana, 2020; Christman et al., 2021). After six weeks of continuous RNAi planarians grew and were fixed and stained with 4’,6-diamidino-2-phenylindole (DAPI) to visualize the anatomy and distribution of testis lobes. As expected, testis lobes were abundant from the along the dorsolateral anatomy of *Luciferase(RNAi)* (Figure 2A), and the same was seen for *Smed-TRPM-l(RNAi)* planarians (Figure 2D). On the other hand, testes lobes were partially or completely missing in regions of *Smed-TRPM-c(RNAi)* and *Smed-TRPM-c;TRPM-l(RNAi)* planarians (Figure 2B-C and 2E). The absence of testis lobes was particularly noticeable at the posterior end of these animals (Figure 2B’-C’, and 2E’), which is a region where testis lobes are normally abundant (Figure 2A’ and 2D’). Beyond distribution of testis lobes, analysis of sperm development by high magnification confocal microscopy revealed full progression of spermatogenesis and spermatozoa accumulation in the innermost region of testis lobes of *Luciferase(RNAi)* and *Smed-TRPM-l(RNAi)* planarians (Figure 2A’’ and 2D’’). In contrast, *Smed-TRPM-c(RNAi)* and *Smed-TRPM-c;TRPM-l(RNAi)* planarians had testis lobes with disruptions at different stages of spermatogenesis. These included loss of spermatozoa (Figure 2B’’), loss of spermatids and/or spermatocytes (Figure 2E’’), and reduced spermatogonia (Figure 2C’’).

**Figure 2.**
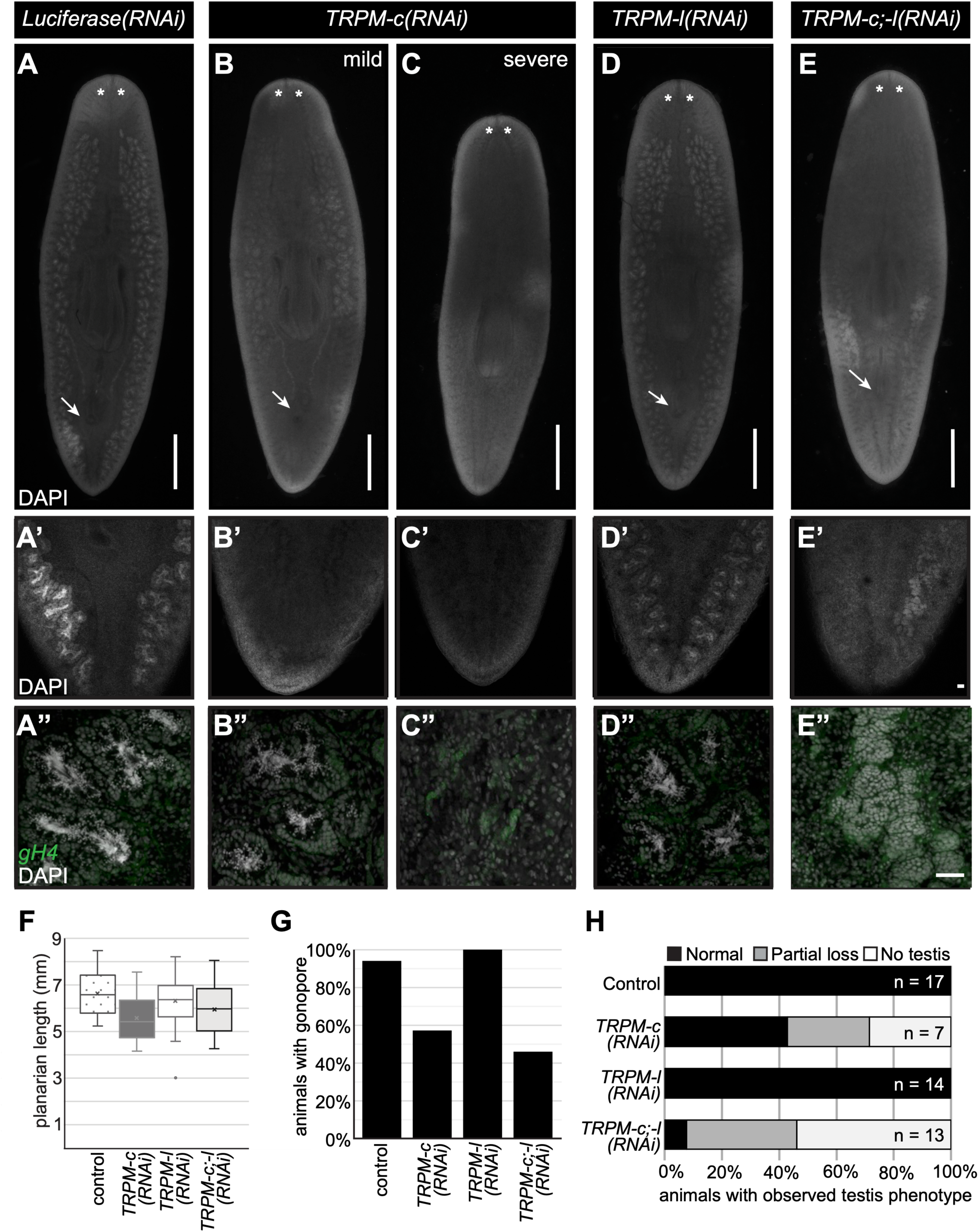
Decreased testis development is observed upon *Smed-TRPM-c* knockdown and *Smed-TRPM-c;TRPM-l* double knockdown. **(A-E)** DAPI signals from whole-mount staining of *Luciferase(RNAi)* (A-A”) and *Smed-TRPM-l(RNAi)* (D-D”) show normal distribution of testis lobes along the dorsolateral anatomy of sexual planarians (A, D). Accumulation of spermatozoa inside testis lobes can be observed in these same samples by confocal microscopy under 10X (A’, D’) and 20X (A”, D”) objectives. Parallel analysis of *Smed-TRPM-c(RNAi)* and *Smed-TRPM-c;TRPM-l(RNAi)* (*TRPM-c;-l(RNAi)*) show partial (B; E) to complete (C) loss of testis lobes and/or testis lobes producing spermatozoa (B”-C”, and E”). Germline stem cells and spermatogonia in testes are shown using *germinal histone H4* (*gH4)* fluorescence *in situ* hybridization signal (green in A”-E”). Development of testis lobes observed in *Luciferase(RNAi)* and *Smed-TRPM-l(RNAi)* posterior regions (A’ and D’) was particularly reduced in *Smed-TRPM-c(RNAi)* and *Smed-TRPM-c;TRPM-l(RNAi)* samples (B’-C’, and E’). **(F)** Box and whisker plot showing average planarian length post-fixation. Mean is marked by an “X”. **(G)** Percentage of animals in each RNAi group displaying a fully developed copulatory apparatus at end of RNAi treatment. **(H)** Quantification of testis development phenotypes (as in A-E) in animals of size comparable to sexually mature controls illustrates percentage of samples with normal testis development (black bar), as well as partial (gray bar) and complete (white bar) absence of testis in each group. Asterisks (*) indicate position of eyes and arrows indicate position of copulatory organ in (A-E). Scale bars = 1 mm (A-E) and 50 μm (E’ and E”).

Sexual maturation and development of reproductive structures are related to growth and somatic integrity (Kobayashi et al., 1999; Wang et al., 2007). Therefore, animal size and development of somatic reproductive structures were analyzed in *Smed-TRPM-c*, *Smed-TRPM-l*, and *Smed-TRPM-c;TRPM-l* knockdowns. *Smed-TRPM-c(RNAi)* displayed reduced average length when compared to the *Luciferase(RNAi)* control and *Smed-TRPM-c(RNAi)* groups (Figure 2F). Additionally, about half of the *Smed-TRPM-c(RNAi)* as well as *Smed-TRPM-c;TRPM-l(RNAi)* planarians failed to develop copulatory structures (Figure 2G), indicating compromised sexual maturation. To assess whether the reduced testis phenotype in *Smed-TRPM-c(RNAi)* resulted from size deficiencies, sperm development was compared between animals of equivalent size or larger than the smallest sexually mature planarian from the *Luciferase(RNAi)* group (i.e. exhibiting fully developed copulatory structures). Amongst these groups, *Smed-TRPM-c(RNAi)* planarians still displayed spermatogenesis defects not observed in the control or *Smed-TRPM-l(RNAi)* groups, and knockdown of *Smed-TRPM-l* in the *Smed-TRPM-c;TRPM-l(RNAi)* group seemed to exacerbate the loss of testis and sperm development (Figure 2H). These results suggest that *Smed-TRPM-c* is required for testis development and/or spermatogenesis directly, and not as a consequence of growth defects.

### Smed-TRPM-c has domains present in mammalian TRPMs and is predicted to form a homotetrameric transmembrane channel

Given that disruption of *Smed-TRPM-c* expression alone resulted in phenotypic observations comparable to *Smed-TRPM-c;TRPM-l* double knockdown, further studies were carried out focusing on *Smed-TRPM-c*. The full cDNA sequence for *Smed-TRPM-c* was obtained by 3’ and 5’ rapid amplification of cDNA ends (RACE), followed by direct amplification, cloning, and sequencing of *Smed-TRPM-c* open reading frames. BLASTP analysis of the protein sequence coded in the longest identified open reading frame (5,241 nts), identified TRPM3 as the closest human homolog to this planarian gene (Figure 3A; E-value = 4E-166), which corroborates with the analysis of reference cDNA contigs for *Smed-TRPM-c* deposited on PlanMine (Supplementary Table S1).

**Figure 3.**
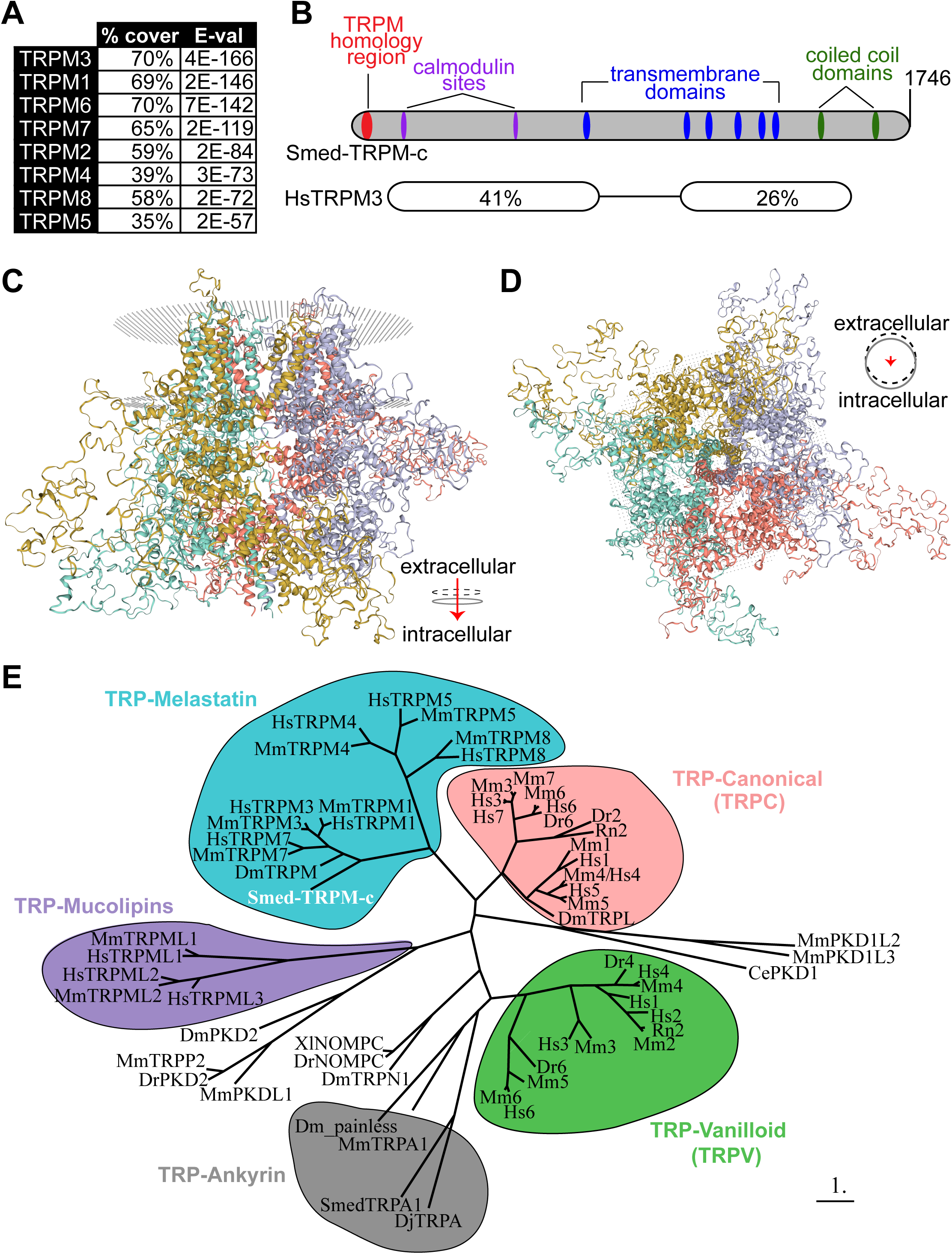
Predicted protein structure and phylogenetic analysis of Smed-TRPM-c. **(A)** The top eight human matches in BLASTP analysis against Smed-PRTM-c are TRPM proteins. **(B)** Architecture of Smed-TRPM-c illustrated using a bar diagram that includes predictions of transmembrane (blue) and coiled coil (green) domains identified by SMART (Simple Modular Architecture Research Tool; Letunic et al., 2020), as well as a TRP-Melastatin (TRPM) specific homology region (Grimm et al., 2003; red), and two calmodulin binding sites identified by the Calmodulin Target Database (http://calcium.uhnres.utoronto.ca/ctdb/ctdb/home.html; magenta). Regions of sequence conservation identified by pairwise BLASTP between Smed-TRPM-c and human TRPM3 (isoform 1; NP_066003.3), including percent identity and corresponding E-values. **(C-D)** The predicted quaternary structure of Smed-TRPM-c as a transmembrane homotetramer according to SWISS-MODEL structural modeling (Waterhouse et al., 2018; Bertoni et al., 2017) based on homology to mouse TRPM7 (Duan et al., 2018). The complex of four subunits is shown in its frontal view (C) and intra to extra cellular view (D). Each of four subunits are shown with different color and the phospholipid bilayer is represented by gray dots. **(E)** Phylogenetic tree using maximum-likelihood principle depicts closer association of Smed-TRPM-c (red bold font) with TRPM from *Drosophila melanogaster* (Dm) and mammalian TRPM3/7/1 orthologs, than with other characterized TRPs. Groups of TRPs belonging to the melastatin (blue), canonical (pink), vanilloid (green), ankyrin (gray), and mucolipins (purple) subfamilies are grouped in bubbles. Abbreviated names showing only first letter of genus and species names (e.g., “Hs” for *Homo sapiens*) were utilized for TRPC and TRPV genes. Scale bar represents 1 substitution per amino acid position.

Smed-TRPM-c contains long regions of conserved sequence that include several predicted structural features present in mammalian TRPMs (Figure 3B). These structural features include the N-terminal TRPM Homology Region (MHR) that is characteristic of members of the TRPM subfamily (Grimm et al., 2003; Wagner et al., 2008; Wang et al., 2018; Winker et al., 2017; Supplementary Figure 1), six transmembrane domains (Supplementary Figure 2), two calmodulin binding sites found in human TRPM3 (Holakovska et al., 2012), and two C-terminal coiled-coil domains (Figure 3B). The MHR of Smed-TRPM-c and human TRPMs are 26-37% identical (Supplementary Figure S1). Additionally, the serine residue at the 1107 position of mouse TRPM7, which is important in modulation of enzyme activity in response to phosphatidylinositol 4,5-bisphosphate levels (Zhelay et al., 2018), is conserved in Smed-TRPM-c (Supplementary Figure S2). Analysis of quaternary structure based on protein homology modeling (SWISS-MODEL; Waterhouse et al., 2018; Bertoni et al., 2017) predicted a homotetrameric transmembrane channel as the top formation for Smed-TRPM-c (Figure 3C and 3D). Each unit in the complex is predicted to contribute to the central pore of the channel (Figure 3D). To infer the phylogenetic relationship between Smed-TRPM-c and TRPs characterized in other experimental models, a phylogeny tree based on maximum-likelihood (Dereeper et al., 2008) was generated (Figure 3E). In this analysis, Smed-TRPM-c groups with a subset of TRPMs that includes mammalian TRPM1, 3, and 7, as well as TRPM from *Drosophila*, which are expressed in spermatogenic cells in mammals (Li et al., 2010) and mediate calcium influx during egg activation in *Drosophila* (Hu et al., 2019), respectively. Altogether, these analyses indicate that Smed-TRPM-c shares sequence and structural features that are typical of mammalian TRPMs and close sequence conservation with those expressed in germ cells of other animals.

### *Smed-TRPM-c* is preferentially expressed in developing sperm and required to maintain sperm production

Expression of *Smed-TRPM-c* was analyzed using riboprobes generated from full-length ORF clones, which confirmed preferential expression in testis lobes (Figure 4A and 4A’; n = 8/8 planarians) and validated WMISH results observed while using partial sequence (Figure 1D). Interestingly, expression of *Smed-TRPM-c* was also detected in the ovaries in a fraction of our samples when using full length riboprobes (Figure 4B and 4B’; n = 2/8 planarians). Non-specific signal from WMISH generated from parallel analyses using a riboprobes against firefly *Luciferase* as negative control was minimal (Figure 4C), confirming the specificity of *Smed-TRPM-c* signals. Expression of *Smed-TRPM-c* was not conclusively detected outside of the gonads, although faint signals were observed in regions of the gut and head (Figure 4A and 4B). However, single cell RNAseq (scRNA-seq) data available for asexual planarians (Plass et al., 2018; Fincher et al., 2018), failed to support the notion that *Smed-TRPM-c* is expressed in the gut or in neurons (Supplementary Figure S3). In fact, there is no corroboration regarding detection of expression of *Smed-TRPM-c* in any somatic cell type between different scRNA-seq studies (Plass et al., 2018; Fincher et al., 2018; Zeng et al., 2018), although one indicated comparably higher expression in glia than is other somatic tissues (Plass et al., 2018; Supplementary Figure S3). These results indicate that *Smed-TRPM-c* is preferentially expressed in the testes.

**Figure 4.**
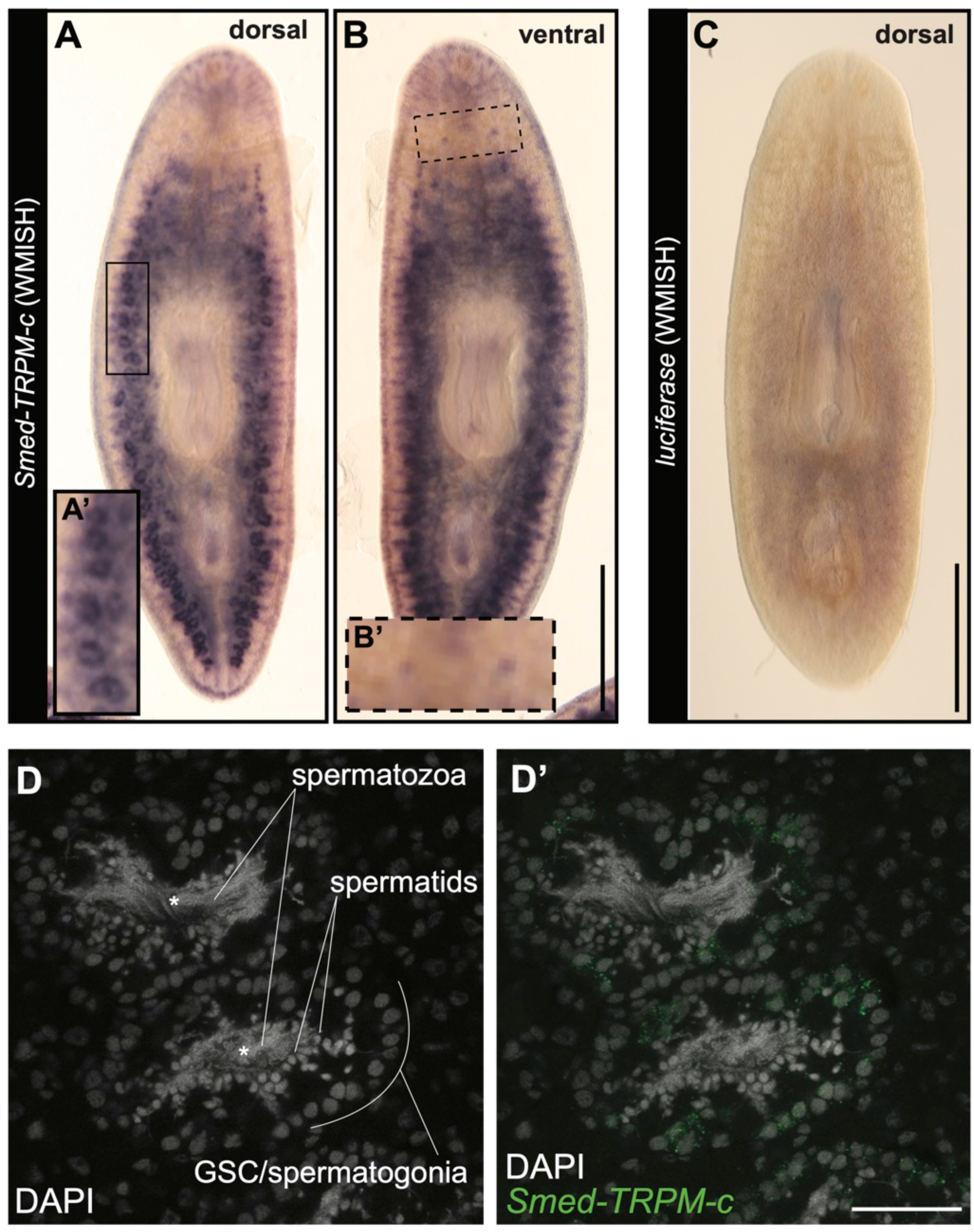
*Smed-TRPM-c* is preferentially expressed in testes. **(A-C)** Whole-mount *in situ* hybridization analysis using full-length ORF probes in sexual planarians shows enriched expression of *Smed-TRPM-c* in testis lobes of sexual planarians (n = 8/8; A) and in the ventrally located ovaries (n = 2/8; B). Background signal generated from negative control *luciferase* riboprobes is shown in (C). Insets show 2-fold magnified views of testis lobes (A’; full frame inset) and ovaries (B’; dashed frame). **(D)** Confocal section displaying the signal detection of *Smed-TRPM-c* fluorescence *in situ* hybridization (green; D’) in the outer cellular region of testis lobes. DAPI stain of nuclear DNA (gray; D and D’) show changes in cellular morphology during the progressive differentiation of spermatogenic cells from the outer to inner region of each lobe. Asterisks mark location of the innermost region of each testis lobe. Scale bar = 1 mm in A-C, and 50 μm in D’.

Fluorescent *in situ* hybridization (FISH) was performed to reveal the cellular distribution of *Smed-TRPM-c* expression in testes in further detail. Planarian testis lobes are primarily populated by germ cells undergoing spermatogenesis (Wang et al., 2007; Newmark et al., 2008; Collins et al., 2010). Spermatogonial stem cells proliferate to form cysts that differentiate progressively as their location changes from periphery to center of each testis lobe (*i.e.*, germline stem cells/spermatogonia in the outer region, and spermatids/elongated spermatozoa in the innermost region of the lobe; Figure 4D). *Smed-TRPM-c* expression was detected most abundantly in the periphery of the testis lobe, which is where male germline stem cells and spermatogonia are located (Figure 4D’). These results indicate that *Smed-TRPM-c* is preferentially expressed during early stages of spermatogenesis.

RNAi experiments performed using dsRNA corresponding to full-length *Smed-TRPM-c* ORF were performed to verify the phenotype observed upon knockdown of this gene using partial sequence. Groups of juvenile *S. mediterranea* were fed dsRNA mixed with liver twice per week. A positive control for RNAi efficacy (*Smed-prohormone convertase 2; pc2(RNAi)*) and a negative control (*Luciferase(RNAi)*) were included for comparison. After six weeks of RNAi, samples of comparable size were stained with DAPI and analyzed by fluorescence stereo and confocal microscopy. DAPI staining revealed normal testis morphology and distribution in all of the *Luciferase(RNAi)* samples (n = 12; Figure 5, A-A’ and C). As was expected from previous studies (Collins et al., 2010), animals from the *pc2(RNAi)* group lacked testes and showed progressively less motility (Figure 5C; not shown), indicating that the RNAi regimen was effective. Interestingly, about half of the animals in the *Smed-TRPM-c(RNAi)* group (n = 23) displayed fewer and underdeveloped testes in comparison to control samples (Figure 5, B-B’ and C). In contrast, analysis of ovaries did not reveal noticeable changes in morphology between *Luciferase(RNAi)* and *Smed-TRPM-c(RNAi)* (Figure 5, A’’ and B’’), and quantification of the number of oocytes present per ovary did noy reveal statistically significant a difference in *Smed-TRPM-c(RNAi)* and control planarians (Figure 5D; mean 12.2 oocytes/ovary in *luciferase(RNAi)* vs. 9.6 oocytes/ovary *Smed-TRPM-c(RNAi)*; unpaired two-tailed student’s *t-* test, p > 0.05). These results indicate that *Smed-TRPM-c* is primarily required to support normal sperm development in planarians.

**Figure 5.**
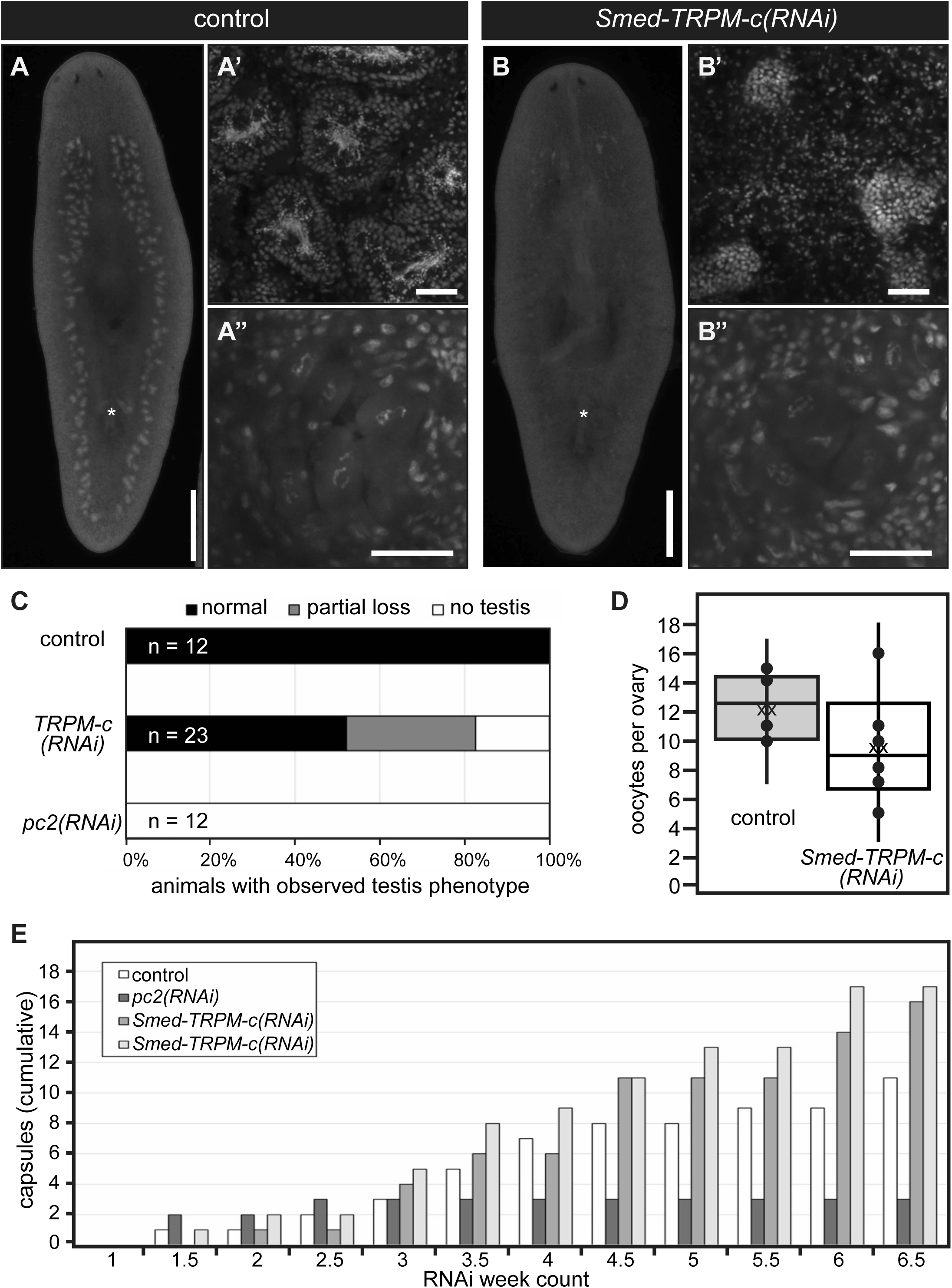
*Smed-TRPM-c* supports sperm development. **(A-B)** Analysis of DAPI-stained samples by whole-mount fluorescence stereomicroscopy reveals testis lobe distribution and presence of copulatory apparatus (asterisks) in *Luciferase(RNAi)* (control; A) and *Smed-TRPM-c(RNAi)* (B) planarians. Analyses of same samples by single-plane confocal microscopy reveal normal sperm development in *Luciferase(RNAi)* planarians **(A’)** and sperm development disruption in *Smed-TRPM-c(RNAi)* **(B’)**, whereas development of oocytes seems comparable **(A” and B”)**. Scale bars = 1 mm (A and B) and 50 μm (A′-B′ and A”-B”). **(C-D)** Quantification of gonad development phenotypes (as in A-B) revealed normal testes distribution (black bar; C) in all control samples, and partial (gray bar) or complete loss (white bar) of productive testes in almost half of all *Smed-TRPM-c(RNAi)* planarians (n = 23). Complete loss of testes was observed in all *pc2(RNAi)* planarians (n = 12). **(D)** Box and whisker plot show a slight decrease in the number of oocytes per ovary in *Smed-TRPM-c(RNAi)* samples (white) when compared to *luciferase(RNAi)* (gray). This difference was not statistically significant (*t-*test, p > 0.05). Dots represent individual data points, “xx” marks the mean. **(E)** Analysis of cumulative egg capsule production as a readout of functional somatic reproductive structures reveals continuous production of capsules in *Luciferase(RNAi)* (control, white bars) and two groups of *Smed-TRPM-c(RNAi)* planarians (lighter shades of gray), whereas *pc2(RNAi)* planarians (dark gray) ceased producing capsules 2.5 weeks into RNAi treatments.

To further check for pleiotropic defects related to maintenance of sexual maturation, production of egg capsules was analyzed in sexually-mature *Smed-TRMP-c(RNAi)* planarians. Planarians, like most flatworms, produce ectolecithal eggs that lack the nutritional components necessary for early development (Gremigni, 1983). Instead, the embryo is encapsulated with yolk cells produced by yolk glands (vitellaria) that develop at the end of sexual maturation using modifications to oogenic genetic programs (Hoshi et al., 2003; Rouhana et al., 2017; Issigonis et al., 2022). Importantly, production and deposition of egg capsules occur independently of presence or absence of functional gametes (Steiner et al., 2016). To test the requirement of *Smed-TRPM-c* in vitellaria maintenance and function, sexually mature planarians were subjected to six weeks of RNAi treatments and the number of egg capsules deposited was recorded after each dsRNA feeding as an indicator of vitellaria presence and function. The number of capsules deposited by *Luciferase(RNAi)* control planarians accumulated continuously over the course of the RNAi treatment (Figure 5E) as previously observed (Rouhana et al., 2017; Steiner et al., 2016). In contrast, *pc2(RNAi)* planarians produced capsules only during the first two weeks of treatment (Figure 5E), which was expected given that *pc2* function is required for maintenance of sexual maturation (Collins et al., 2010). *Smed-TRPM-c(RNAi)* planarians produced egg capsules continuously over the course of the experiment, reaching even higher quantities than the *luciferase(RNAi)* group (Figure 5E), and indicating that *Smed-TRPM-c* is not required for yolk cell development, maintenance, or function.

In addition to testing for capsule formation and deposition, changes in behavior were investigated to assess potential roles of *Smed-TRPM-c* in the brain or the possibility of non-specific knockdown of TRP paralogs with functions in *S. mediterranea* sensory behaviors. Chemotaxis response was indirectly tested by tracking the fraction of planarians that ate during RNAi feedings, for which no differences were observed (data not shown). To test for changes in behavioral response to temperature, an arena composed of two cold (17°C) and two hot (30°C) quadrants was manufactured using a design similar to that described by Arenas et al. (2017). *Luciferase(RNAi)* spent most of their time in the cold quadrants and avoided hot temperature quadrants (Supplementary Figure 4), as expected from findings by Arenas et al. (2017). No significant difference was observed in the time spent in cold and hot quadrants between *Smed-TRPM-c(RNAi)* and control groups (Supplementary Figure 4), supporting the notion that *Smed-TRPM-c* is specifically required for sperm development and not involved in neuronal functions that regulate behavior.

### *Smed-TRPM-c* supports development and/or maintenance of *nanos*(+) presumptive germline stem cells in asexual planarians

The phenotype observed from RNAi analyses in sexual planarians indicated that *Smed-TRPM-c* supports early stages of spermatogenesis. To further investigate this hypothesis, expression of *Smed-TRPM-c* was disrupted in asexual planarians. An asexual strain of *S. mediterranea* that reproduces exclusively through fission and regeneration contains a chromosomal translocation assumed to impede sexual maturation (Newmark and Sanchez Alvarado, 2002; Newmark et al., 2008; Guo et al., 2022). However, planarians in this asexual strain still contain presumptive germline stem cells located in the dorsolateral position equivalent to where testes develop in sexual planarians (Wang et al., 2007). Groups of asexual *S. mediterranea* were subjected to six-weeks of *Luciferase* or *Smed-TRPM-c* dsRNA feedings. Upon completion of dsRNA feedings, the presence of presumptive germline stem cell clusters was visualized using *nanos(+)* marker expression by WMISH. At the end of the 6-week RNAi treatment clusters of *nanos(+)* cells were quantified in both *Luciferase(RNAi)* control planarians (Figure 6A-A’) and *Smed-TRPM-c(RNAi)* planarians (Figure 6B-B’). Plotting the abundance of *nanos(+)* clusters against the length of each animal revealed that the number of *nanos(+)* clusters was fewer in *Smed-TRPM-c* knockdowns than in control animals of comparable size (Figure 6C). Statistical analysis using two-tailed Student’s *t*-test indicated that the difference in number of clusters per millimeter of animal length between *Smed-TRPM-c(RNAi)* and control planarians is significant (p < 0.05). This reduction in relative number of *nanos(+)* clusters corroborates with the loss of testes lobes observed in *Smed-TRPM-c(RNAi)* sexual planarians (Figure 2B, 2C, 2H, 5B and 5C). Given these results, we propose that *Smed-TRPM-c* contributes to spermatogenesis through regulation of establishment, maintenance, and/or expansion of germline stem cells.

**Figure 6.**
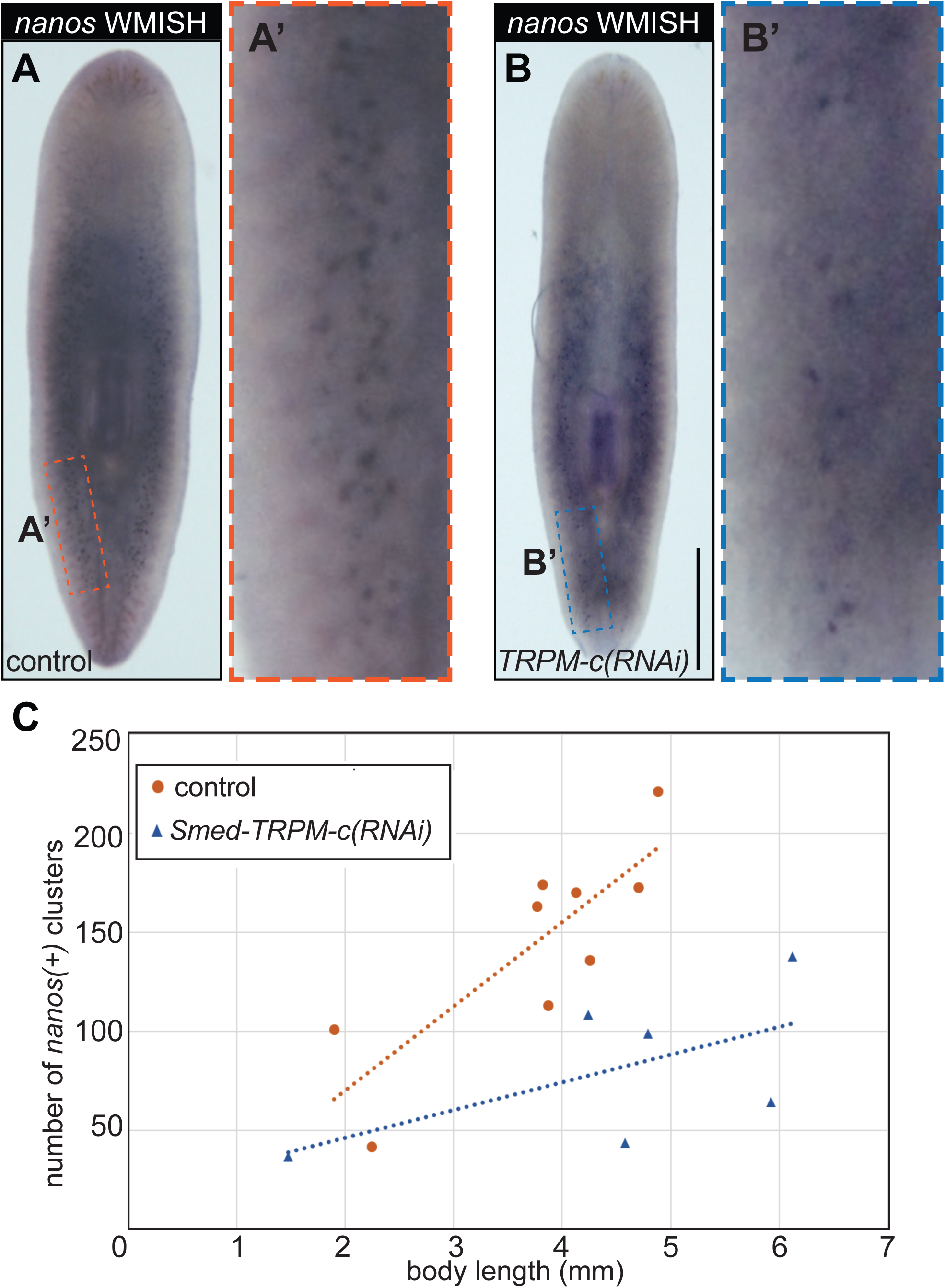
*Smed-TRPM-c* is required for maintenance of presumptive germline stem cells in asexual planarians. **(A-B)** Distribution of *nanos(+)* germline stem cell clusters in control *Luciferase(RNAi)* (A) and *Smed-TRPM-c(RNAi)* (B) asexual planarians are displayed in bright field microscopy images of samples subjected to whole-mount *in situ* hybridization. Insets in **(A’ and B’)** show 5-fold magnified view. Scale bar = 1 mm. **(C)** Quantitative analysis *nanos(+)* cluster number as detected in (A-B; y-axis) is plotted in relation to planarian body length (x-axis).

## METHODS

### Planarian cultures

A laboratory line of sexual *S. mediterranea* (Zayas et al., 2005) was used for all experiments except for those indicated as specifically using asexual animals, in which case specimens from the CIW4 clonal laboratory strain were used (Newmark and Sánchez Alvarado, 2000). Asexual animals were maintained in 1X Montjui□c salts at 21°C. Sexual animals were maintained in 0.75X Montjui□c salts at 18°C. Planarians were maintained in the dark except during weekly or biweekly feedings with beef calf liver, which were done on bench tops at room temperature. Animals were starved for at least one week prior to experimentation or fixation.

### Identification of TRPM homologs and generation of riboprobes for whole-mount *in situ* hybridization

TRPM homologs in *S. mediterranea* were identified from transcriptomes of sexual (dd_Smes_v1 prefix) and asexual (dd_Smed_v6 prefix) strains deposited in PlanMine (Rozanski et al., 2018) using human TRPM3 (NCBI GenBank ID NP_066003.3) as input in TBLASTN searches. Redundant records were identified by pairwise BLASTN comparisons and removed. GeneArt Strings DNA fragments (ThermoFisher, Waltham, MA) were synthesized for each *S. mediterranea* TRPM homolog to contain ∼500 bps of reference contig sequence flanked by SP6 (sense) and T3 (antisense) promoter sequences (Supplementary File 1). These custom-ordered DNA fragments were amplified by PCR using primers corresponding of SP6 and T3 promoter sequences that included an additional T7 promoter sequence at their 5’-ends (5′-GAATTTAATACGACTCACTATAGGGCGATTTAGGTGACACTATAGA-AGAGAAC-3′ and 5′-GAATTTAATACGACTCACTATAGGGCGAATTAACCCTCACTAAAGGGAAC-3′, respectively) as described in Counts and Rouhana (2017). PCR fragments were purified using DNA Clean & Concentrator-5 kits as per the manufacturer (Zymo Research, Irvine, CA), eluted in 20 μL or RNase-free water, and used as templates for synthesis of riboprobes labeled with Digoxigenin-11-UTP (Roche, distributed by MilliporeSigma, Burlington, MA) using T3 RNA polymerase (Promega, Madison, WI).

### Cloning of *Smed-TRPM-c* cDNA

Full-length mRNA sequence for *Smed-TRPM-c* was predicted from matching contigs (dd_Smed_v6_11377_0_1, dd_Smes_v1_41098_1_4, uc_Smed_v2_27847, others) identified in in PlanMine (Rozanski et al., 2018) and primers for 5’ and 3’ RACE [5’-AGGAAGCAATGATTTACCGGGAGAAAG-3’ and 5’-CAAGTGGTCAGTAGAGCGCATTGACTAC-3’] were designed based on this prediction. Full-length *Smed-TRPM-c* ORF was amplified using Long PCR Master Mix (Promega, Madison, WI) from sexual *S. mediterranea* cDNA synthesized using oligo(dT) and N_6_ primers. The oligo primers used for amplification of *Smed-TRPM-c* ORF were 5’-GTCACAGTGGCATTGTAGCCAATCCCTC-3’ and 5’-ATTGGCGAGTAAATCGCTTTGCATTGC-3’. The forward primer is position immediately upstream from the first 19 nucleotides of the ORF (5’-ATGAAAAAATCTAAAAAAA-3’) according to records on PlanMine. Primers were not designed to include this region due to rich A/T content. Amplicons were ligated into the vector pGEM-T (Promega, Madison, WI) and verified by Sanger sequencing.

### Bioinformatic analysis of protein structure

The transmembrane and coiled coil domains of the predicted *Smed-TRPM-c* full-length ORF were identified using the Normal SMART program (Letunic et al., 2014). The starting motif of the TRPM Homology Region (MHR) was derived from Grimm et al. (2003). The putative calmodulin binding sites of Smed-TRPM-c were identified with the Calmodulin Target Database (http://calcium.uhnres.utoronto.ca/ctdb/ctdb/home.html). N-terminus TRPM homology region analysis was performed by alignment of TRPM Homology Region of different TRPM channels (Grimm et al., 2003) using ClustalW 2.1 (Thompson et al., 1994). The multiple alignment was then input into BoxShade (https://github.com/mdbaron42/pyBoxshade/releases) using the default settings. Predictive models of three-dimensional Smed-TRPM-c structure were obtained using SWISS-MODEL structural modeling (https://swissmodel.expasy.org/; Waterhouse et al., 2018; Bertoni et al., 2017) based on homology to mouse TRMP7 (Duan et al., 2018) and using default settings.

### Smed-TRPM-c phylogenetic analysis

TRP protein sequences were obtained from analyses published in Inoue et al. (2014) and Arenas et al. (2017). The predicted amino acid sequence of Smed-TRPM-c was aligned with those of TRP proteins from these studies using ClustalW and used as input to generate a phylogenetic tree using PhyML and TreeDyn in phylogeny.fr (Dereeper et al., 2008). Default settings were used across programs.

### Whole-mount *in situ* hybridization

Whole-mount *in situ* hybridization was performed as described by Pearson et al. (2009) with modifications in fixation steps following King and Newmark (2013). Large sexually-mature animals (∼1.0 cm or larger) were used, therefore the N-acetylcysteine (NAC) treatment, as well as the initial fixation step and the Proteinase K treatment were prolonged. Planarians were placed in a solution of 10% NAC in PBS for 11 minutes with gentle agitation, then fixed in 4% formaldehyde in PBS containing 0.3% Triton X-100 (PBSTx) and rocked for 1 hour at 4°C. The animals were then gradually dehydrated in methanol and kept at −20°C overnight. The following day, the samples were gradually rehydrated in PBSTx and bleached in formamide bleaching solution (King and Newmark, 2013) for 2 hours under a bright light. After bleaching, the samples were rinsed in PBSTx and placed in a Proteinase K solution containing 10μg/ml of Proteinase K (ThermoFisher, Waltham, MA) in PBSTx with 0.1% (w/v) SDS for 12 minutes with gentle agitation. The samples were post-fixed for 10 minutes with 4% formaldehyde in PBSTx. The steps for hybridization and post-hybridization washes were performed as indicated by Pearson et al. (2009). For colorimetric *in situ* hybridization, samples were incubated in blocking solution containing 5% horse serum in TNTx (consisting of 0.1M Tris pH 7.5, 0.15M NaCl, and 0.3% Triton X-100) for at least 2 hours. Samples subjected to colorimetric signal detection were incubated in anti-Digoxygenin-AP antibody solution (1:4000 dilution; Roche Diagnostics, Mannheim, Germany) overnight while rocking at 4°C. The post-antibody incubation washes with TNTx and signal development were performed as in Pearson et al. (2009) for colorimetric signal detection. After development, the samples were mounted in 80% glycerol/20% PBS and imaged using a Zeiss V.16 SteREO microscope equipped with a Canon EOS Rebel T3 camera. For fluorescent *in situ* hybridization, samples were incubated in blocking solution containing 5% horse serum and 1% Western Blocking Reagent (Roche Diagnostics, Mannheim, Germany) in TNTx for at least 2 hours. Samples used in FISH analyses were incubated in anti-DIG-POD antibody solution (1:2000 dilution; Roche Diagnostics, Mannheim, Germany) rocking overnight at 4°C. The FISH samples were developed using FAM tyramide solution as described by King and Newmark (2013), washed, and mounted in 80% glycerol/20% PBS and imaged using a Nikon C2+ confocal microscope with NIS Elements C software. Initial observations of distribution of expression of TRPM homologs were performed using riboprobes generated from GeneArt DNA fragments, and *Smed-TRPM-c* expression was validated in colorimetric and FISH analyses using riboprobes generated from full-length ORF clones.

### Disruption of *Smed-TRPM-c* expression by RNA-interference (RNAi)

Templates for *in vitro* transcription of dsRNA generated by PCR from GeneArt fragments using the aforementioned primers (see “generation of riboprobes for whole-mount *in situ* hybridization, above) or from DNA cloned into pGEM-T (Promega, Madison, WI) using the following primers: 5’-GAATTTAATACGACTCACTATAGGGCGCCAAGCTATTTAGGTGACACTATAGAATACTC-3’ and 5’-GAATTAATTAACCCTCACTAAAGGGAGAATTTAATACGACTCACTATAGGGCGAATTGG-3’). PCR products were purified using DNA Clean & Concentrator-5 columns (Zymo Research, Irvine, CA), eluted in 20μl of RNase-free water and used as templates for *in vitro* transcription. DsRNA was synthesized using T7 RNA polymerase as described by Rouhana et al. (2013). Groups of 6 juvenile sexual planarians (lacking gonopores), mature sexual planarians actively laying capsules, or 10 asexual planarians 0.5-0.7 mm length, were fed to satiation with a liver solution containing approximately 100 ng/μl of gene-specific dsRNA twice per week for at least 4 weeks. As a negative control, one group of planarians was fed firefly *Luciferase* dsRNA which is not expressed in the planarian genome and has no effect on their sexual development nor behavior. To assess RNAi efficacy in real-time, some experiments included one group of planarians fed with *Smed-pc2* dsRNA which is required for testes maintenance and normal planarian behavior (Reddien et al., 2005; Collins et al., 2010).

### Analysis of testis distribution and sperm development

A week following completion of the last RNAi feeding, planarian testis distribution and anatomy were assessed by DAPI staining. The samples were fixed and bleached as described above for *in situ* hybridization (minus methanol dehydration and rehydration steps). After bleaching, the samples were washed in PBSTx twice and then incubated in DAPI (1:1000 dilution of 1mg/ml DAPI stock solution in PBSTx; ACROS Organics, Morris, NJ) overnight while rocking at 4°C. After incubating overnight, the samples were washed 4 times with PBSTx and then mounted on slides with 4:1 glycerol:PBS and imaged under UV light with a Zeiss V.16 SteREO microscope equipped with a Canon EOS Rebel T3 camera (for low magnification) or a Nikon C2+ confocal microscope using a 10X or 20X objective and running NIS Elements C software (for high magnification).

### Analysis of ovarian anatomy, oocyte development, and egg capsule deposition

To assess possible disruptions to oogenesis, the number of oocytes per ovary of *Smed-TRPM-c* knockdown planarians were compared to *luciferase* knockdown planarians. Juvenile sexual planarians fed gene-specific dsRNA for at least 4 weeks were stained with DAPI a week following the conclusion of the RNAi feeding regimen. The number of oocytes were counted using Z-stacks on a Nikon C2+ confocal microscope with NIS Elements C software. The ovaries of five *luciferase(RNAi)* and five *Smed-TRPM-c(RNAi)* planarians were examined. Each count of oocytes was plotted as an individual point in a box and whisker plot. Statistical analysis was performed using unpaired two-tailed Student’s *t*-test. To assess egg capsule deposition, groups of 6 sexually mature planarians (1.0 – 1.5 cm in length) were used per group. The animals were fed dsRNA as described above. The cumulative number of egg capsules deposited was recorded after each dsRNA feeding.

### Analysis of presumptive germline stem cell distribution in asexual planarians

To assess distribution of presumptive germline stem cells in asexual planarians, groups of asexual *S. mediterranea* were subjected to dsRNA feedings for six weeks, fixed a week following the last feeding, and processed for colorimetric *in situ* hybridization using a *nanos* riboprobe (Wang et al., 2007). After *in situ* hybridization, each animal was imaged with a Zeiss V.16 SteREO microscope equipped with a Canon EOS Rebel T3 camera and the length of each animal was determined by drawing a 3-point arc from the center of the head to the pharynx and to the center of the tail of each planarian image using Adobe Illustrator. The images were each assigned a random number and subjected to triple-blind analysis. The number of *nanos(+)* clusters were counted by three individuals, averaged, and plotted against the length of the animal. Statistical analysis was completed using two-tailed unequal variance Student’s *t*-test on the ratios between the number of *nanos* clusters and the length of the animal.

### Thermotaxis assay

Thermotactic behavior of asexual planarians was tested with an aluminum temperature plate based on the design used by Arenas et al. (2017) (Supplementary Figure S4). The aluminum temperature plate was controlled by a simple variable 12-volt power supply. The power supply was regulated by two temperature controllers (Elitech, Milpitas, CA). The temperature controllers fed four Peltier plates in diagonal quadrants. Each pair of Peltier plates were either powered by positive or negative 12-volt direct current. Positive current produced warmer above-ambient temperatures while the negative current produced below-ambient temperatures.

To assemble the apparatus, thermal silver paste was used to adhere the anodized aluminum plate to the Peltier plates. The aluminum plate was treated with a hydrophobic water-proofing spray while leaving an uncoated circle of diameter 8.6 centimeters in the center. Two temperature sensors were adhered to the perimeter of two adjacent quadrants of the untreated circle. The heat sink was adhered to the bottom of the Peltier plates using the same thermal paste. A small computer fan was attached to the bottom of the heat sink to assist with heat dispersion.

The untreated circle was filled with Montjui□c salts to a height of approximately 3 mm, forming a bubble of water for the samples to move through. Groups of 10 asexual planarians were subjected to six weeks of RNAi treatment and then starved for one week prior to experimentation. Four *Luciferase(RNAi)* planarians and five *Smed-TRPM-c(RNAi)* planarians were fed pureed beef liver mixed with colored chalk shavings (Hagoromo Co., Ltd., Kasugai, Japan) as per Hattori et al. (2018) prior to being placed on the temperature plate to increase the visibility of the planarians during the assay. The nine planarians were placed in the center of the circle together and their movements were recorded for 10 minutes. Heat map images were taken with a thermal camera (Hti-Xintai, Dongguan, Guangdong, China) at five-minute intervals. The percentage of time spent in cold quadrants for each individual planarian was calculated from the video recording and plotted on Excel. Statistical analysis was performed using two-tailed unequal variance Student’s *t*-test.

## DISCUSSION

Our research revealed 21 potential TRPM subfamily members in the planarian *S. mediterranea.* Expression of two of these was detected in testes, while some of the others were expressed in the gut, head tip, and subsets of neurons (Figure 1). *Smed-TRPM-c* expression is enriched in testes (Figure 1 and Figure 4), which parallels expression of *TRPM3* in mouse testis (Jang et al., 2012) and in rat spermatogenic cells (Li et al., 2010). Expression in the ovaries was only observed in a subset of animals possibly due to incomplete development of the ovaries, which occurs towards the end of sexual maturation. *Smed-TRPM-c* is required to maintain normal distribution and development of active testes in the dorsolateral anatomy of sexual planarians (Figure 2 and Figure 5) and presumptive germ cell clusters in asexual planarians (Figure 6). We hypothesize that *Smed-TRPM-c* contributes to spermatogenesis via cell-autonomous mechanisms based on detection of expression of this gene in the outer layer of germ cells in testis lobes, which is where spermatogonial stem cells and spermatogonia reside (Figure 4D-D’). We also hypothesize that *Smed-TRPM-c* works to regulate sperm development in the earliest steps of spermatogenesis (*i.e.*, establishment of germline stem cell proliferation and/or maintenance), based on the observation that presumptive spermatogonial stem cell clusters are less abundant upon reduction of *Smed-TRPM-c* expression in asexual *S. mediterranea* (Figure 6). These results place Smed-TRPM-c as a sensor of either cellular or environmental cues that drive sperm development in *S. mediterranea*.

Members of the TRPM channel subfamily are known to be activated by cold (TRPM8; McKemy et al., 2002) and hot temperatures (TRPM3; Vriens et al., 2011), as well as by chemical ligands such as the steroid hormone pregnenolone sulphate and spermine (reviewed by Huang et al., 2020). The stimuli that activate Smed-TRPM-c remain unknown. However, our preferred hypothesis is that Smed-TRPM-c is responsive to temperature and regulates cell-autonomous mechanisms in the male germline. This is based on three indirect observations: 1) A member of the TRPM subfamily that displays preferential expression during spermatogenesis in mammals protects germ cells from damages induced by cold-shock (TRPM8; Borowiec et al., 2016); 2) Smed-TRPM-c shares highest sequence conservation of TRPM3 amongst mammalian orthologs, and TRPM3 is responsive to heat; 3) sexual reproduction and development in planarians are heavily regulated by temperature (see below); and 4) we didn’t detect any sperm development defects upon knockdown of orthologs of genes involved in pregnenolone sulphate metabolism (data not shown).

Planarians inhabit freshwater ecosystems all over the world, with Antarctica and islands where colonization has not occurred being possible exceptions (Vila-Farré and Rink, 2018). Most species in the genus *Schmidtea* are primarily distributed in northern Europe, while *S. mediterranea* has been mainly reported to inhabit southern European regions such as Tunisia, Italy, and Catalonia (Lázaro et al., 2011). Exclusively fissiparous populations (*i.e.*, asexually reproducing through fission and regeneration) of *S. mediterranea* and other planarian species are not uncommon. However, sexual reproduction by hermaphroditism is the predominant and ancestral mechanism for reproduction in planarians. Development, size, and activity of planarian reproductive structures in the wild follow seasonal patterns, and some planarian species even switch between sexual and fissiparous reproductive strategies (Dahm, 1958; Ball and Reynoldson, 1981). Temperature has long-been an abiotic factor of interest in modulation of planarian reproductive strategies. For example, *S. mediterranea* testes, ovaries, and the copulatory apparatus shrink and eventually disappear in water temperatures above 20°C in (Harrath et al., 2004). However, the environmental and molecular factors that regulate this sexual maturation and reproduction in the wild remain to be uncovered.

Some evidence suggests that temperature works to tune sexual maturity in planarians through independent tissue-specific pathways, rather than through a master regulator. For example, *Schmidtea (Dugesia) lugubris* exhibits regressed testes at temperatures below 5°C, but these grow and become fertile when transferred to 10°C (Reynoldson et al., 1965). In contrast, *S. lugubris* ovaries are populated with oocytes even at temperatures below 5°C, indicating that testes and ovaries do not follow the same temperature restrictions. In another example, *Polycelis tenuis* produces capsules below 5°C, indicating that vitellaria are present and functional, but these capsules are sterile. Given that gametes are not required for capsule production in planarians (Steiner et al., 2016), the observation that *P. tenuis* produces capsules at low temperatures indicates that vitellaria are fully developed and functional, while their sterility suggests that sperm, ova, or both are absent or dysfunctional. In contrast, the planarian species *Dendrocoelum lacteum* produces fertile capsules at temperatures as low as 1°C to 5°C, indicating that functional oocytes, sperm, and vitellaria are present at these low temperatures. The stimulus or stimuli that activate Smed-TRPM-c remain to be found. However, *Smed-TRPM-c* does not seem to function as a “master switch” that modulates reproductive maturation across tissues. Instead, *Smed-TRPM-c* sems to serve primarily in the testis and promote specification, proliferation, and/or maintenance of spermatogonial germline stem cells. Tissue-specific functions of sensory receptors like *Smed-TRPM-c* provide a mechanism by which planarians can evolve reproductive system strategies in ways that maximize fitness in the different freshwater ecosystems that they inhabit.

## ACKNOWLEDGEMENTS

The authors would like to thank Donovan Christman M.S. for his assistance in the triple-blind analysis of *nanos*(+) clusters in sexual planarians, Jacob Rissler for help with RNAi experiments, as well as Ashot Kozak and Jananie Rockwood for attempts to test planarian TRPM activity *in vitro*. This work was supported by a grant from The Eunice Kennedy Shriver National Institute of Child Health and Human Development of the National Institutes of Health (R15HD082754) to LR.

## REFERENCES

Arenas OM, Zaharieva EE, Para A, Vásquez-Doorman C, Petersen CP, Gallio M. Activation of planarian TRPA1 by reactive oxygen species reveals a conserved mechanism for animal nociception. Nat Neurosci. 2017 Dec;20(12):1686–1693. doi: 10.1038/s41593-017-0005-0. Epub 2017 Oct 16. PMID: 29184198; PMCID: PMC5856474.

Ball IR, Reynoldson TB (1981) British planarians. Synopsis of the British fauna. N. 19. Cambridge University Press, Cambridge

Bertoni M, Kiefer F, Biasini M, Bordoli L, Schwede T. Modeling protein quaternary structure of homo- and hetero-oligomers beyond binary interactions by homology. Sci Rep. 2017 Sep 5;7(1):10480. doi: 10.1038/s41598-017-09654-8. PMID: 28874689; PMCID: PMC5585393.

Borowiec AS, Sion B, Chalmel F, D Rolland A, Lemonnier L, De Clerck T, Bokhobza A, Derouiche S, Dewailly E, Slomianny C, Mauduit C, Benahmed M, Roudbaraki M, Jégou B, Prevarskaya N, Bidaux G. Cold/menthol TRPM8 receptors initiate the cold-shock response and protect germ cells from cold-shock-induced oxidation. FASEB J. 2016 Sep;30(9):3155–70. doi: 10.1096/fj.201600257R. Epub 2016 Jun 17. PMID: 27317670; PMCID: PMC5001517.

Brandl H, Moon H, Vila-Farré M, Liu SY, Henry I, Rink JC. PlanMine--a mineable resource of planarian biology and biodiversity. Nucleic Acids Res. 2016 Jan 4;44(D1):D764–73. doi: 10.1093/nar/gkv1148. Epub 2015 Nov 17. PMID: 26578570; PMCID: PMC4702831.

Christman DA, Curry HN, Rouhana L. Heterotrimeric Kinesin II is required for flagellar assembly and elongation of nuclear morphology during spermiogenesis in Schmidtea mediterranea. Dev Biol. 2021 Sep;477:191–204. doi: 10.1016/j.ydbio.2021.05.018. Epub 2021 Jun 4. PMID: 34090925; PMCID: PMC8277772.

Chmura HE, Williams CT. A cross-taxonomic perspective on the integration of temperature cues in vertebrate seasonal neuroendocrine pathways. Horm Behav. 2022 Aug;144:105215. doi: 10.1016/j.yhbeh.2022.105215. Epub 2022 Jun 7. PMID: 35687987.

Collins JJ 3rd, Hou X, Romanova EV, Lambrus BG, Miller CM, Saberi A, Sweedler JV, Newmark PA. Genome-wide analyses reveal a role for peptide hormones in planarian germline development. PLoS Biol. 2010 Oct 12;8(10):e1000509. doi: 10.1371/journal.pbio.1000509. Erratum in: PLoS Biol. 2015 Aug;13(8):e1002234. PMID: 20967238; PMCID: PMC2953531.

Counts JT, Hester TM, Rouhana L. Genetic expansion of chaperonin-containing TCP-1 (CCT/TRiC) complex subunits yields testis-specific isoforms required for spermatogenesis in planarian flatworms. Mol Reprod Dev. 2017 Dec;84(12):1271–1284. doi: 10.1002/mrd.22925. Epub 2017 Nov 10. PMID: 29095551; PMCID: PMC5760344.

Dahm AG (1958) Taxonomy and ecology of five species groups in the family Planariidae (Turbellaria Tricladida Paludicola). Nya Litografen, Malm.

De Blas GA, Darszon A, Ocampo AY, Serrano CJ, Castellano LE, Hernández-González EO, Chirinos M, Larrea F, Beltrán C, Treviño CL. TRPM8, a versatile channel in human sperm. PLoS One. 2009 Jun 30;4(6):e6095. doi: 10.1371/journal.pone.0006095. PMID: 19582168; PMCID: PMC2705237.

Dereeper A, Guignon V, Blanc G, Audic S, Buffet S, Chevenet F, Dufayard JF, Guindon S, Lefort V, Lescot M, Claverie JM, Gascuel O. Phylogeny.fr: robust phylogenetic analysis for the non-specialist. Nucleic Acids Res. 2008 Jul 1;36(Web Server issue):W465–9. doi: 10.1093/nar/gkn180. Epub 2008 Apr 19. PMID: 18424797; PMCID: PMC2447785.

Diver MM, Lin King JV, Julius D, Cheng Y. Sensory TRP Channels in Three Dimensions. Annu Rev Biochem. 2022 Jun 21;91:629–649. doi: 10.1146/annurev-biochem-032620-105738. Epub 2022 Mar 14. PMID: 35287474; PMCID: PMC9233036.

Duan J, Li Z, Li J, Hulse RE, Santa-Cruz A, Valinsky WC, Abiria SA, Krapivinsky G, Zhang J, Clapham DE. Structure of the mammalian TRPM7, a magnesium channel required during embryonic development. Proc Natl Acad Sci U S A. 2018 Aug 28;115(35):E8201–E8210. doi: 10.1073/pnas.1810719115. Epub 2018 Aug 14. PMID: 30108148; PMCID: PMC6126765.

Fincher CT, Wurtzel O, de Hoog T, Kravarik KM, Reddien PW. Cell type transcriptome atlas for the planarian *Schmidtea mediterranea*. Science. 2018 May 25;360(6391):eaaq1736. doi: 10.1126/science.aaq1736. Epub 2018 Apr 19. PMID: 29674431; PMCID: PMC6563842.

Gremigni V. (1983). Platyhelminthes-Turbellaria. Reproductive Biology of Invertebrates, Vol. 1. Oogenesis, oviposition, and oosorption. New York, NY: Wiley, 67–107.

Grimm C, Kraft R, Sauerbruch S, Schultz G, Harteneck C. Molecular and functional characterization of the melastatin-related cation channel TRPM3. J Biol Chem. 2003 Jun 13;278(24):21493–501. doi: 10.1074/jbc.M300945200. Epub 2003 Apr 2. PMID: 12672799.

Harrath AH, Charni M, Sluys R, Zghal F, Tekaya S. Ecology and distribution of the freshwater planarian *Schmidtea mediterranea* in Tunisia. Italian Journal of Zoology. 2004; 71:3, 233–236, doi: 10.1080/11250000409356577

Hattori M, Miyamoto M, Hosoda K, Umesono Y. Usefulness of multiple chalk-based food colorings for inducing better gene silencing by feeding RNA interference in planarians. Dev Growth Differ. 2018 Jan;60(1):76–81. doi: 10.1111/dgd.12413. Epub 2017 Dec 21. PMID: 29266402.

Holakovska B, Grycova L, Jirku M, Sulc M, Bumba L, Teisinger J. Calmodulin and S100A1 protein interact with N terminus of TRPM3 channel. J Biol Chem. 2012 May 11;287(20):16645–55. doi: 10.1074/jbc.M112.350686. Epub 2012 Mar 27. PMID: 22451665; PMCID: PMC3351314.

Hu Q, Wolfner MF. The *Drosophila* Trpm channel mediates calcium influx during egg activation. Proc Natl Acad Sci U S A. 2019 Sep 17;116(38):18994–19000. doi: 10.1073/pnas.1906967116. Epub 2019 Aug 19. PMID: 31427540; PMCID: PMC6754564.

Huang Y, Fliegert R, Guse AH, Lü W, Du J. A structural overview of the ion channels of the TRPM family. Cell Calcium. 2020 Jan;85:102111. doi: 10.1016/j.ceca.2019.102111. Epub 2019 Nov 24. PMID: 31812825; PMCID: PMC7050466.

Hyman LH. (1951). North American triclad Turbellaria. XII. Synopsis of the known species of fresh-water planarians of North America. Transactions of the American Microscopical Society, 70(2), 154–167.

Inoue T, Yamashita T, Agata K. Thermosensory signaling by TRPM is processed by brain serotonergic neurons to produce planarian thermotaxis. J Neurosci. 2014 Nov 19;34(47):15701–14. doi: 10.1523/JNEUROSCI.5379-13.2014. PMID: 25411498; PMCID: PMC6608440.

Issigonis M, Browder KL, Chen R, Collins JJ, Newmark PA. A niche-derived non-ribosomal peptide triggers planarian sexual development. bioRxiv [Preprint]. 2023 Dec 6:2023.12.06.570471. doi: 10.1101/2023.12.06.570471. PMID: 38106172; PMCID: PMC10723454.

Issigonis M, Newmark PA. From worm to germ: Germ cell development and regeneration in planarians. Curr Top Dev Biol. 2019;135:127–153. doi: 10.1016/bs.ctdb.2019.04.001. Epub 2019 May 2. PMID: 31155357.

Issigonis M, Redkar AB, Rozario T, Khan UW, Mejia-Sanchez R, Lapan SW, Reddien PW, Newmark PA. A Krüppel-like factor is required for development and regeneration of germline and yolk cells from somatic stem cells in planarians. PLoS Biol. 2022 Jul 15;20(7):e3001472. doi: 10.1371/journal.pbio.3001472. PMID: 35839223; PMCID: PMC9286257.

Jang Y, Lee Y, Kim SM, Yang YD, Jung J, Oh U. Quantitative analysis of TRP channel genes in mouse organs. Arch Pharm Res. 2012 Oct;35(10):1823–30. doi: 10.1007/s12272-012-1016-8. Epub 2012 Nov 9. PMID: 23139135.

Kashio M, Tominaga M. TRP channels in thermosensation. Curr Opin Neurobiol. 2022 Aug;75:102591. doi: 10.1016/j.conb.2022.102591. Epub 2022 Jun 18. PMID: 35728275.

Kenk, R. (1935). STUDIES ON VIRGINIAN TRICLADS. Journal of the Elisha Mitchell Scientific Society, 51(1), 79–125.

King RS, Newmark PA. In situ hybridization protocol for enhanced detection of gene expression in the planarian Schmidtea mediterranea. BMC Dev Biol. 2013 Mar 12;13:8. doi: 10.1186/1471-213X-13-8. PMID: 23497040; PMCID: PMC3610298.

Kobayashi, K., R. Koyanagi, M. Matsumoto, J. P. Cabrera, and M. Hoshi. 1999. Switching from asexual to sexual reproduction in the planarian *Dugesia ryukyuensis*: Bioassay system and basic description of sexualizing process. Zool. Sci. 16:291–298.

Kobayashi K, Matsumoto M, Hoshi M. [Switching mechanism from asexual to sexual reproduction in planarians]. Tanpakushitsu Kakusan Koso. 2004 Feb;49(2):102–7. Japanese. PMID: 14969100.

Lázaro EM, Harrath AH, Stocchino GA, Pala M, Baguñà J, Riutort M. Schmidtea mediterranea phylogeography: an old species surviving on a few Mediterranean islands? BMC Evol Biol. 2011 Sep 26;11:274. doi: 10.1186/1471-2148-11-274. PMID: 21943163; PMCID: PMC3203090.

Lee N, Chen J, Sun L, Wu S, Gray KR, Rich A, Huang M, Lin JH, Feder JN, Janovitz EB, Levesque PC, Blanar MA. Expression and characterization of human transient receptor potential melastatin 3 (hTRPM3). J Biol Chem. 2003 Jun 6;278(23):20890–7. doi: 10.1074/jbc.M211232200. Epub 2003 Apr 2. PMID: 12672827.

Lesko SL, Rouhana L. Dynein assembly factor with WD repeat domains 1 (DAW1) is required for the function of motile cilia in the planarian Schmidtea mediterranea. Dev Growth Differ. 2020 Aug;62(6):423–437. doi: 10.1111/dgd.12669. Epub 2020 Jun 1. PMID: 32359074; PMCID: PMC7483408.

Letunic I, Khedkar S, Bork P. SMART: recent updates, new developments and status in 2020. Nucleic Acids Res. 2021 Jan 8;49(D1):D458–D460. doi: 10.1093/nar/gkaa937. PMID: 33104802; PMCID: PMC7778883.

Li S, Wang X, Ye H, Gao W, Pu X, Yang Z. Distribution profiles of transient receptor potential melastatin- and vanilloid-related channels in rat spermatogenic cells and sperm. Mol Biol Rep. 2010 Mar;37(3):1287–93. doi: 10.1007/s11033-009-9503-9. Epub 2009 Mar 26. PMID: 19322679.

Magley RA, Rouhana L. Tau tubulin kinase is required for spermatogenesis and development of motile cilia in planarian flatworms. Mol Biol Cell. 2019 Aug 1;30(17):2155–2170. doi: 10.1091/mbc.E18-10-0663. Epub 2019 May 29. PMID: 31141462; PMCID: PMC6743461.

Martínez-López P, Treviño CL, de la Vega-Beltrán JL, De Blas G, Monroy E, Beltrán C, Orta G, Gibbs GM, O’Bryan MK, Darszon A. TRPM8 in mouse sperm detects temperature changes and may influence the acrosome reaction. J Cell Physiol. 2011 Jun;226(6):1620–31. doi: 10.1002/jcp.22493. PMID: 21413020.

McKemy DD, Neuhausser WM, Julius D. Identification of a cold receptor reveals a general role for TRP channels in thermosensation. Nature. 2002 Mar 7;416(6876):52-8. doi: 10.1038/nature719. Epub 2002 Feb 10. PMID: 11882888.

Nakagawa H, Sekii K, Maezawa T, Kitamura M, Miyashita S, Abukawa M, Matsumoto M, Kobayashi K. A comprehensive comparison of sex-inducing activity in asexual worms of the planarian *Dugesia ryukyuensis*: the crucial sex-inducing substance appears to be present in yolk glands in Tricladida. Zoological Lett. 2018 Jun 12;4:14. doi: 10.1186/s40851-018-0096-9. Erratum in: Zoological Lett. 2018 Aug 31;4:25. PMID: 29942643; PMCID: PMC5996458.

Newmark PA, Sánchez Alvarado A. Bromodeoxyuridine specifically labels the regenerative stem cells of planarians. Dev Biol. 2000 Apr 15;220(2):142–53. doi: 10.1006/dbio.2000.9645. PMID: 10753506.

Newmark PA, Sánchez Alvarado A. Not your father’s planarian: a classic model enters the era of functional genomics. Nat Rev Genet. 2002 Mar;3(3):210–9. doi: 10.1038/nrg759. PMID: 11972158.

Newmark PA, Wang Y, Chong T. Germ cell specification and regeneration in planarians. Cold Spring Harb Symp Quant Biol. 2008;73:573–81. doi: 10.1101/sqb.2008.73.022. Epub 2008 Nov 6. PMID: 19022767.

Pearson BJ, Eisenhoffer GT, Gurley KA, Rink JC, Miller DE, Sánchez Alvarado A. Formaldehyde-based whole-mount in situ hybridization method for planarians. Dev Dyn. 2009 Feb;238(2):443–50. doi: 10.1002/dvdy.21849. PMID: 19161223; PMCID: PMC2640425.

Plass M, Solana J, Wolf FA, Ayoub S, Misios A, Glažar P, Obermayer B, Theis FJ, Kocks C, Rajewsky N. Cell type atlas and lineage tree of a whole complex animal by single-cell transcriptomics. Science. 2018 May 25;360(6391):eaaq1723. doi: 10.1126/science.aaq1723. Epub 2018 Apr 19. PMID: 29674432.

Reynoldson TB, Young JO, Taylor MC. The effect of temperature on the life-cycle of four species of lake-dwelling triclads. J. Anim. Ecol. 1965, 34: 23–43 10.2307/2367.

Rouhana L, Tasaki J, Saberi A, Newmark PA. Genetic dissection of the planarian reproductive system through characterization of Schmidtea mediterranea CPEB homologs. Dev Biol. 2017 Jun 1;426(1):43–55. doi: 10.1016/j.ydbio.2017.04.008. Epub 2017 Apr 21. PMID: 28434803; PMCID: PMC5544531.

Rouhana L, Weiss JA, Forsthoefel DJ, Lee H, King RS, Inoue T, Shibata N, Agata K, Newmark PA. RNA interference by feeding in vitro-synthesized double-stranded RNA to planarians: methodology and dynamics. Dev Dyn. 2013 Jun;242(6):718–30. doi: 10.1002/dvdy.23950. Epub 2013 Apr 1. PMID: 23441014; PMCID: PMC3909682.

Rozanski A, Moon H, Brandl H, Martín-Durán JM, Grohme MA, Hüttner K, Bartscherer K, Henry I, Rink JC. PlanMine 3.0-improvements to a mineable resource of flatworm biology and biodiversity. Nucleic Acids Res. 2019 Jan 8;47(D1):D812–D820. doi: 10.1093/nar/gky1070. PMID: 30496475; PMCID: PMC6324014.

Thompson JD, Higgins DG, Gibson TJ. CLUSTAL W: improving the sensitivity of progressive multiple sequence alignment through sequence weighting, position-specific gap penalties and weight matrix choice. Nucleic Acids Res. 1994 Nov 11;22(22):4673–80. doi: 10.1093/nar/22.22.4673. PMID: 7984417; PMCID: PMC308517.

Vila-Farré M, C Rink J. The Ecology of Freshwater Planarians. Methods Mol Biol. 2018;1774:173–205. doi: 10.1007/978-1-4939-7802-1_3. PMID: 29916156.

Vila-Farré M, Rozanski A, Ivanković M, Cleland J, Brand JN, Thalen F, Grohme MA, von Kannen S, Grosbusch AL, Vu HT, Prieto CE, Carbayo F, Egger B, Bleidorn C, Rasko JEJ, Rink JC. Evolutionary dynamics of whole-body regeneration across planarian flatworms. Nat Ecol Evol. 2023 Dec;7(12):2108–2124. doi: 10.1038/s41559-023-02221-7. Epub 2023 Oct 19. PMID: 37857891; PMCID: PMC10697840.

Vriens J, Owsianik G, Hofmann T, Philipp SE, Stab J, Chen X, Benoit M, Xue F, Janssens A, Kerselaers S, Oberwinkler J, Vennekens R, Gudermann T, Nilius B, Voets T. TRPM3 is a nociceptor channel involved in the detection of noxious heat. Neuron. 2011 May 12;70(3):482–94. doi: 10.1016/j.neuron.2011.02.051. PMID: 21555074.

Wagner TF, Loch S, Lambert S, Straub I, Mannebach S, Mathar I, Düfer M, Lis A, Flockerzi V, Philipp SE, Oberwinkler J. Transient receptor potential M3 channels are ionotropic steroid receptors in pancreatic beta cells. Nat Cell Biol. 2008 Dec;10(12):1421–30. doi: 10.1038/ncb1801. Epub 2008 Nov 2. PMID: 18978782.

Wang L, Fu TM, Zhou Y, Xia S, Greka A, Wu H. Structures and gating mechanism of human TRPM2. Science. 2018 Dec 21;362(6421):eaav4809. doi: 10.1126/science.aav4809. Epub 2018 Nov 22. PMID: 30467180; PMCID: PMC6459600.

Wang Y, Stary JM, Wilhelm JE, Newmark PA. A functional genomic screen in planarians identifies novel regulators of germ cell development. Genes Dev. 2010 Sep 15;24(18):2081–92. doi: 10.1101/gad.1951010. PMID: 20844018; PMCID: PMC2939369.

Wang Y, Zayas RM, Guo T, Newmark PA. nanos function is essential for development and regeneration of planarian germ cells. Proc Natl Acad Sci U S A. 2007 Apr 3;104(14):5901–6. doi: 10.1073/pnas.0609708104. Epub 2007 Mar 21. PMID: 17376870; PMCID: PMC1851589.

Waterhouse A, Bertoni M, Bienert S, Studer G, Tauriello G, Gumienny R, Heer FT, de Beer TAP, Rempfer C, Bordoli L, Lepore R, Schwede T. SWISS-MODEL: homology modelling of protein structures and complexes. Nucleic Acids Res. 2018 Jul 2;46(W1):W296–W303. doi: 10.1093/nar/gky427. PMID: 29788355; PMCID: PMC6030848.

Winkler PA, Huang Y, Sun W, Du J, Lü W. Electron cryo-microscopy structure of a human TRPM4 channel. Nature. 2017 Dec 14;552(7684):200-204. doi: 10.1038/nature24674. Epub 2017 Dec 6. PMID: 29211723.

Zayas RM, Hernández A, Habermann B, Wang Y, Stary JM, Newmark PA. The planarian Schmidtea mediterranea as a model for epigenetic germ cell specification: analysis of ESTs from the hermaphroditic strain. Proc Natl Acad Sci U S A. 2005 Dec 20;102(51):18491–6. doi: 10.1073/pnas.0509507102. Epub 2005 Dec 12. Erratum in: Proc Natl Acad Sci U S A. 2012 Nov 13;109(46):19033. PMID: 16344473; PMCID: PMC1317986.

Zhelay T, Wieczerzak KB, Beesetty P, Alter GM, Matsushita M, Kozak JA. Depletion of plasma membrane-associated phosphoinositides mimics inhibition of TRPM7 channels by cytosolic Mg^2+^, spermine, and pH. J Biol Chem. 2018 Nov 23;293(47):18151–18167. doi: 10.1074/jbc.RA118.004066. Epub 2018 Oct 10. PMID: 30305398; PMCID: PMC6254349.

Zeng A, Li H, Guo L, Gao X, McKinney S, Wang Y, Yu Z, Park J, Semerad C, Ross E, Cheng LC, Davies E, Lei K, Wang W, Perera A, Hall K, Peak A, Box A, Sánchez Alvarado A. Prospectively Isolated Tetraspanin^+^ Neoblasts Are Adult Pluripotent Stem Cells Underlying Planaria Regeneration. Cell. 2018 Jun 14;173(7):1593–1608.e20. doi: 10.1016/j.cell.2018.05.006. PMID: 29906446; PMCID: PMC9359418.

Zheng J. Molecular mechanism of TRP channels. Compr Physiol. 2013 Jan;3(1):221–42. doi: 10.1002/cphy.c120001. PMID: 23720286; PMCID: PMC3775668.

