## Supplementary Material Legends for "Melastatin subfamily Transient Receptor Potential channels support spermatogenesis in planarian flatworms"

**Supplementary File 1. Sequence of GeneArt Strings Gene Synthesis constructs used for initial analyses of *S. mediterranea* TRPM homologs.** Sequence of templates used to generate amplicons for *in vitro* transcription of DIG-labeled riboprobes and dsRNA against *S. mediterranea* TRPMs. Partial *Smed-TRPM* paralog reference contig sequences (lower case font) are flanked by sense SP6 and antisense T3 RNA Polymerase promoter sequences in their 3’- and 5’- ends (capitalized font), respectively.

**Supplementary Table S1. TRPM homologs identified in *Schmidtea mediterranea*.** Twenty-one different TRPM homologs (*TRPM-a1*, *-a2*, and *-b* to *-t*; left column) were identified through TBLASTN searches against *S. mediterranea* transcriptomes (first column on left). Corresponding Smes_v1 and dd_Smed_v6 identifiers are indicated (second and fifth columns, respectively). Top hit amongst human proteins (third column) identified through BLASTX searches using predicted full length cDNA contigs and corresponding E-values (fourth column). Distribution of expression of each TRPM homolog according to scRNA-seq analysis in asexual planarians (Plass et al., 2018; sixth column) and *in situ* hybridization analysis (ISH) in sexual planarians (this work; right column).

**Supplementary Figure 1. Alignment of Human and Smed-TRPM-c TRPM Homology Regions (MHRs).** MHRs of human TRPM channels (Grimm et al., 2003) were aligned using ClustalW 2.1 (Dereeper et al., 2008) and used as input for BoxShade (default settings). Identically conserved positions (black) and positions with changes that conserved similar amino acid properties (gray) in the majority of the sequences analyzed are highlighted.

**Supplementary Figure 2. Alignment of mouse TRPM3, human TRPMs, and Smed-TRPM-c transmembrane regions.** ClustalW 2.1 (Dereeper et al., 2008) alignment of mammalian TRPM homologs with Smed-TRPM-c transmembrane region reveals conserved sequence between transmembrane domains (red arrows) and serine residue (magenta arrow) known to be important in modulation of enzyme activity in response to PIP2 levels (Zhelay et al., 2018). Identically conserved positions (black) and positions with changes that conserved similar amino acid properties (gray) in the majority of the sequences analyzed are highlighted.

**Supplementary Figure 3. Distribution of *Smed-TRPM-c* expression in asexual planarians according to single-cell RNAseq (scRNA-seq) studies.** Images obtained from PlanMine (Rozanski et al., 2018) display enriched detection of *Smed-TRPM-c* in **(A)** glia (Plass et al.,2018) and **(B)** neoblasts (Fincher et al., 2018). **(C)** ScRNA-seq studies focused on planarian stem cells (Zeng et al., 2018) did not detect enrichment of *Smed-TRPM-c* detection in neoblasts.

**Supplementary Figure 4. *Smed-TRPM-c(RNAi)* does not exhibit deficits in planarian thermotactic behavior. (A)** Thermotaxis assay set-up inspired by Arenas et al. (2017) including the temperature display and control panel (green arrow) and the aluminum temperature plate with temperature probes attached (magenta arrow). **(B)** The temperature plate consists of an anodized aluminum plate (1); four Peltier plates (2), two with positive DC electric and two with negative DC electric; an aluminum heat sink for heat dispersion (3); and a fan to increase heat dispersion (4). **(C)** Heat map images of the temperature plate taken with a thermal camera at five and ten minutes into the assay recording (top). Snapshots of the thermotaxis plate under white light at the same timepoints (bottom) show asexual *luciferase(RNAi)* (yellow arrowheads) and *Smed-TRPM-c(RNAi)* (pink arrowheads) planarians in the cold (17°C) quadrants. **(D)** Box and whisker plot showing the range in the percentage of time spent in cold quadrants during the ten-minute experiment of control and *Smed-TRPM-c* knockdown planarians was not statistically significant. Mean is shown with an “x”. Median is shown with a horizontal line. Open circles represent the percentage for each individual planarian.
