## Supplementary Table 1 for "Melastatin subfamily Transient Receptor Potential channels support spermatogenesis in planarian flatworms"

| **Putative TRPM** | **Match Smes_v1 contig** | **Reciprocal BLASTX vs human** | **E-value** | **Match dd_Smed_v6 contig (top match to Smes_v1)** | **Asexual Smed scRNAseq distribution (Plass/Planmine)** | **ISH result** |
| --- | --- | --- | --- | --- | --- | --- |
| **Smed-TRPM-a1**  **(DjTRPMa; E-val = 0)** | dd_Smes_v1_33400_1_1 | TRPM3 | 5.0E-78 | dd_Smed_v6_17857_0_1 | GABA neurons | Peripheral neurons |
| **Smed-TRPM-a2**  **DjTRPMa; E-val = 2.04 × 10^-135^)** | dd_Smes_v1_28579_1_1 | TRPM3 | 5.0E-71 | dd_Smed_v6_26481_0_1 | Goblet cells, GABA neurons | Intestine |
| **Smed-TRPM-b**  **(DjTRPMb; E-val = 0)** | dd_Smes_v1_16865_1_1 | TRPM3 | 8.0E-90 | dd_Smed_v6_9288_0_1 | Secretory 4, pharynx, epidermis, neoblasts | Brain |
| **Smed-TRPM-c** | dd_Smes_v1_41098_1_4 | TRPM3 | 3E-172 | dd_Smed_v6_11377_0_1 | Glia, neoblasts | Testis |
| **Smed-TRPM-d** | dd_Smes_v1_16476_1_2 | TRPM3 | 1.0E-102 | dd_Smed_v6_13119_0_1 | spp-11+ neurons, epidermis, pigment cells | - |
| **Smed-TRPM-e** | dd_Smes_v1_35607_1_1 | TRPM3 | 6.0E-83 | dd_Smed_v6_10029_0_1 | Epidermis | - |
| **Smed-TRPM-f** | dd_Smes_v1_37543_1_1 | TRPM5 | 6.0E-45 | dd_Smed_v6_17981_0_1 | ChAT neurons 1 | Brain, and peripheral neurons |
| **Smed-TRPM-g** | dd_Smes_v1_21604_1_3 | TRPM1 | 1.0E-41 | dd_Smed_v6_21927_0_1 | Secretory 1, secretory 3, protonephridia | - |
| **Smed-TRPM-h** | dd_Smes_v1_41250_2_1 | TRPM2 | 7.0E-84 | dd_Smed_v6_11259_0_1 | npp-18+ neurons, GABA neurons, cav-1+ neurons, secretory 4 | Brain |
| **Smed-TRPM-i** | dd_Smes_v1_30648_1_2 | TRPM2 | 8.0E-69 | dd_Smed_v6_8493_0_1 | otf+ cells 1, 2, glia, aqp+ parenchymal cells, ldlrr-1+ parenchymal cells | - |
| **Smed-TRPM-j** | dd_Smes_v1_33523_1_2 | TRPM3 | 5.0E-61 | dd_Smed_v6_18934_0_6 | Epidermal progenitors, pgrn+ parenchymal cells, phagocytes | Head tip region |
| **Smed-TRPM-k** | dd_Smes_v1_32737_1_3 | TRPM2 | 6.0E-74 | dd_Smed_v6_7784_0_1 | ldlrr-1+, pgrn+, and aqp+ parenchymal cells, phagocytes, glia | - |
| **Smed-TRPM-l** | dd_Smes_v1_89367_1_1 | TRPM8 | 4.0E-31 | dd_Smed_v6_6825_0_1 | pgrn+ parenchymal cells, glia | Intestine and testis |
| **Smed-TRPM-m** | dd_Smes_v1_43220_1_1 | TRPM1 | 6.0E-24 | dd_Smed_v6_10717_0_1 | Epidermal progenitors, epidermis, epidermis DVb, pharynx cell type | Brain |
| **Smed-TRPM-n** | dd_Smes_v1_4986_1_1 | TRPM1 | 5.0E-16 | dd_Smed_v6_13669_0_1 | Epidermal progenitors, epidermis, epidermis DVb, pharynx cell type | Head tip region |
| **Smed-TRPM-o** | dd_Smes_v1_23507_1_1 | TRPM3 | 6.0E-21 | dd_Smed_v6_8382_0_1 | otf+ cells 1, npp-18+ neurons | Intestine |
| **Smed-TRPM-p** | dd_Smes_v1_56329_1_1 | TRPM3 | 1.0E-12 | dd_Smed_v6_15270_0_1 | Epidermal DVb neoblasts, phagocytes, psd+ cells, epidermis, secretory 1 cells | - |
| **Smed-TRPM-q** | dd_Smes_v1_10084_1_1 | TRPM4 | 8.0E-11 | dd_Smed_v6_26251_0_1 | Epidermal DVb | - |
| **Smed-TRPM-r** | dd_Smes_v1_19870_1_4 | TRPM5 | 1.0E-09 | dd_Smed_v6_3620_0_1 | Glia, aqp+ parenchymal cells | Brain and ventral nerve cords |
| **Smed-TRPM-s** | dd_Smes_v1_34165_1_3 | TRPM6 | 2.0E-05 | dd_Smed_v6_10485_0_1 | Phagocytes | Intestine |
| **Smed-TRPM-t** | dd_Smes_v1_68197_1_1 | TRPM3 | 2.0E-06 | dd_Smed_v6_15098_0_1 | Epidermis, goblet cells, protonephridia | - |
