## Supplementary Figure 1 for "Melastatin subfamily Transient Receptor Potential channels support spermatogenesis in planarian flatworms"

```
HTRPM2        59 LSSWIPENIKKKECVYFVESSKLSDAGK---------VVCQCGYTHEQH
HTRPM8        41 LVNFIQANFKKRECVFFTKDSKATEN------------VCKCGYAQSQH
HTRPM3        61 QKSWIERAFYKRECVHIIPSTKDPH-------------RCCCGRLIGQH
HTRPM6         3 QKSWIKGVFDKRECSTIIPSSKNPHRCTPVCQVCQNLIRCYCGRLIGDH
HTRPM7         3 QKSWIESTLTKRECVYIIPSSKDPHRCLPGCQICQQLVRCFCGRLVKQH
HTRPM1       -90 QKSWIEKTFCKRECIFVIPSMKDSN-------------RCCCGQFTNQH
HTRPM4         4 EQSWIPKIFKKKTCTTFIVDSTDPGG-----------TLCQCGRPRTAH
SmedTRPM-c    31 KYNWIDDNILKLECNRFHP--KSNG-------------YCACGRPAEEH
```
