## Supplementary figures and images for "Melastatin subfamily Transient Receptor Potential channels support spermatogenesis in planarian flatworms"

### Supplementary Figure 2

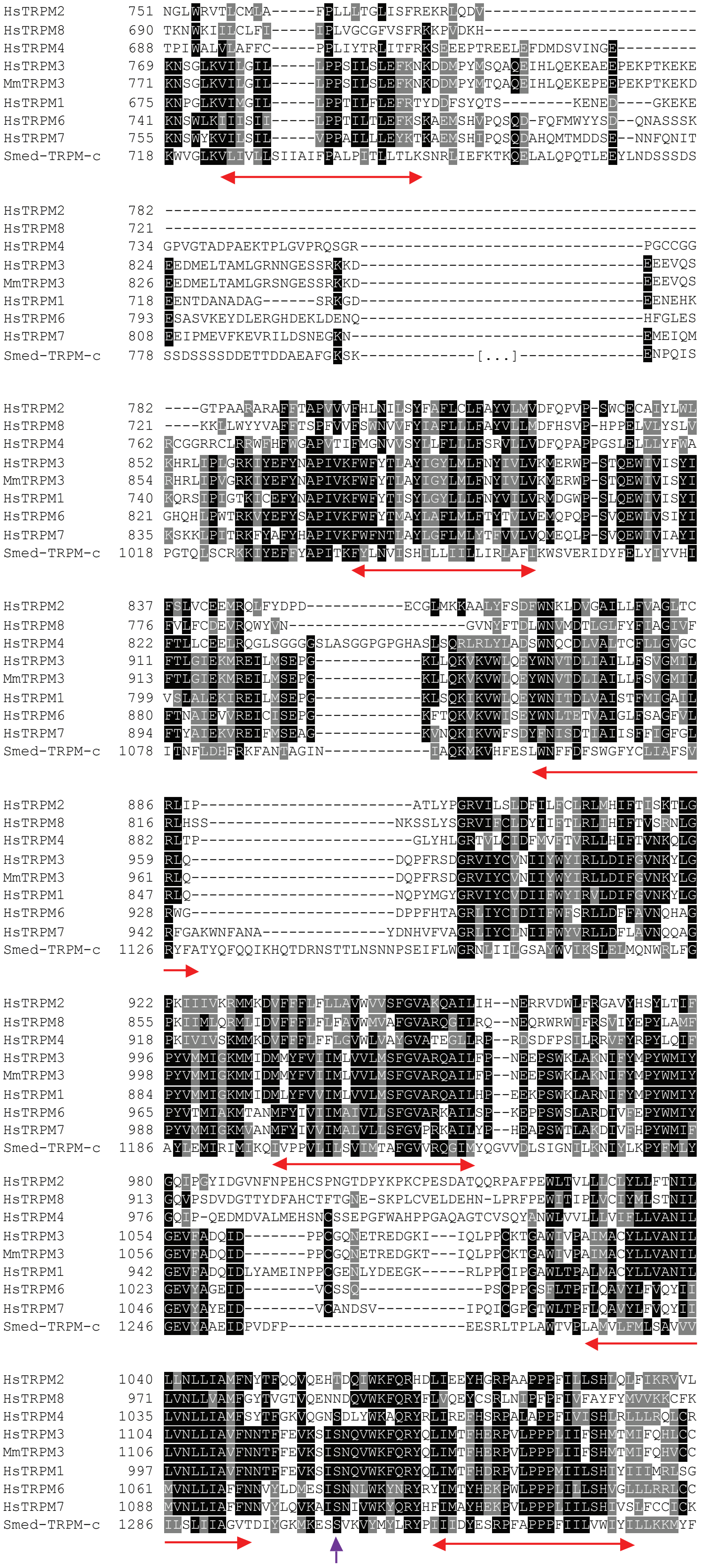

### Supplementary Figure 4

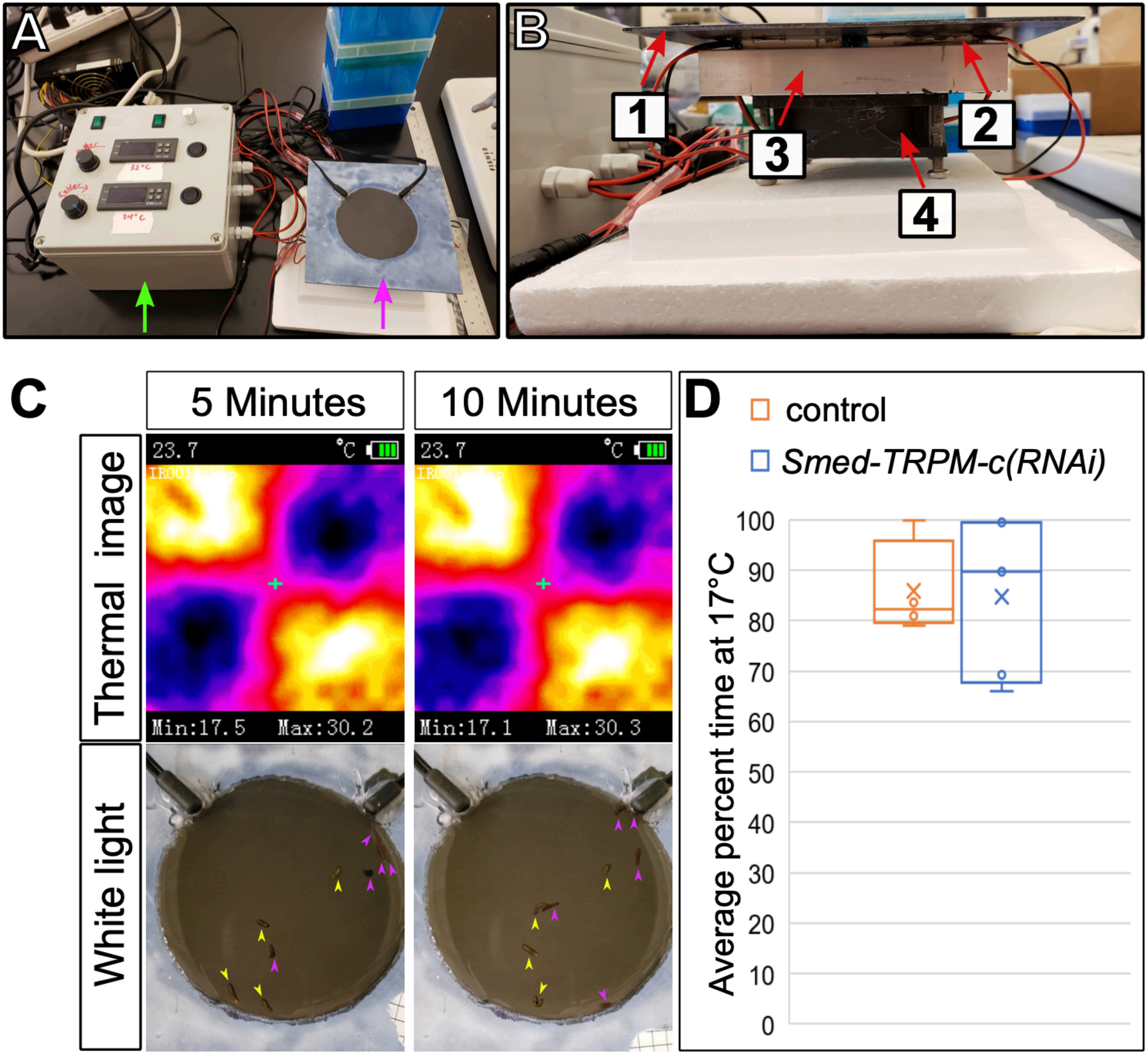
