## Supplementary Figure 3 for "Melastatin subfamily Transient Receptor Potential channels support spermatogenesis in planarian flatworms"

**A** *Smed-TRPM-c* expression according to scRNAseq by (Plass et al., 2018).

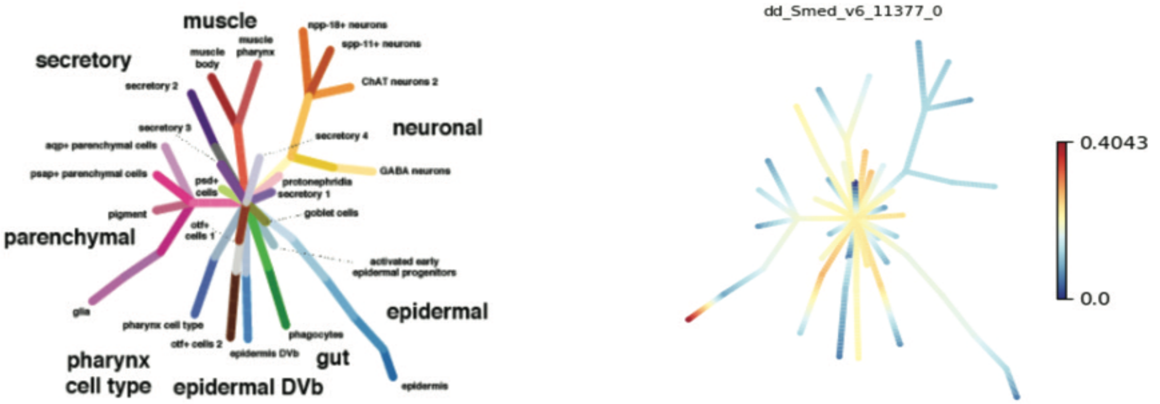


**B** *Smed-TRPM-c* expression according to scRNAseq by (Fincher et al., 2018).

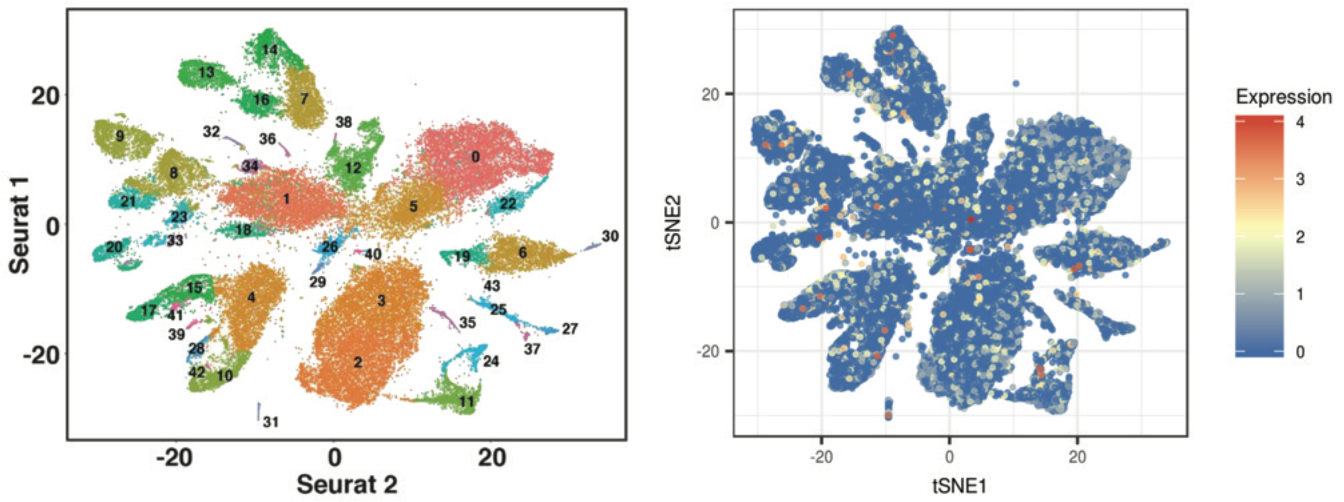


**C** *Smed-TRPM-c* expression according to scRNAseq by (Zeng et al., 2018).

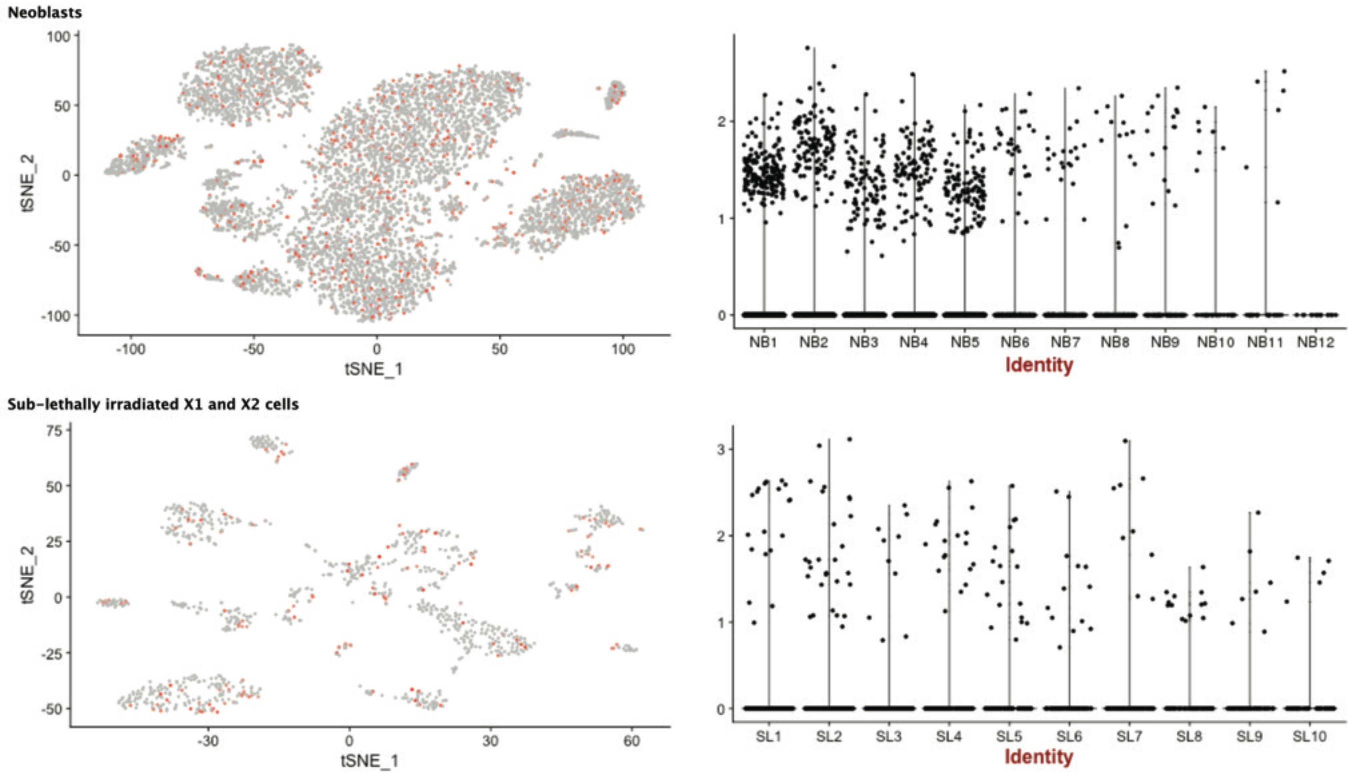
